# Design principles of neuromorphic computing using genetic circuits

**DOI:** 10.64898/2025.12.01.691482

**Authors:** Frank Britto Bisso, Durga Shree, Yinan Zhu, Christian Cuba Samaniego

## Abstract

Cells have evolved to sense a wide range of input combinations and integrate those signals through signaling pathways to produce context-specific responses, such as differentiation, cell-type specification, and patterning. To replicate this information-processing capacity, synthetic biology has developed large-scale circuitry inspired by the fundamental principles of computer science. Within this framework, neuromorphic computing implemented using genetic circuits offers the opportunity to significantly enhance the computational capabilities of single cells. In this work, we establish design principles for implementing neuromorphic computing in living cells by identifying the key feature that enables a chemical reaction network to function as a perceptron: an input-output mapping with a tunable threshold. We demonstrate that four ubiquitous chemical reaction networks, namely molecular sequestration, catalytic degradation, competitive binding, and activation/deactivation cycles, all satisfy this requirement and can be engineered as perceptrons. By layering these perceptrons into multi-layer architectures, we then show how to construct both linear and nonlinear decision boundaries through rational tuning of production rates that encode network weights. As proof of principle, we apply this framework to design neural networks capable of discriminating between healthy and cancer cells based on gene expression data from 19 tissue types. Together, this work formalizes the design principles for engineering genetic circuits as neural networks and establishes a foundation for implementing next-generation cellular computation.

## 1 Introduction

Synthetic biology can be broadly described as operating at three levels: sensing, computation, and actuation. Sensing modules convert extracellular cues into intracellular signals through engineered receptors [1, 2], while actuation modules implement phenotypic responses through the expression of an effector molecule [3, 4]. Computation provides the intermediate layer that interprets the sensed signals and determines which cellular response should be triggered. To engineer this decision-making process in single cells, the biocomputing field has introduced a variety of genetic circuit designs, broadly categorized into two paradigms of computation. Digital computing treats gene expression as effectively “on” or “off”, and uses combinatorial promoters [5, 6, 7, 8], split transcription factors [9, 10] and dimerization reactions [11] to implement logic gates, such as AND, OR, and NOT, as well as digital memory units [12, 13] and arithmetic operations [14]; all producing discrete, all-or-none outputs. Analog computing instead generates outputs that vary continuously with the input levels, extending the repertoire of operations to filters [15, 16, 17], logarithmic sensing [18], gradient detection [19], and oscillations [20, 21].

With these tools at hand, many real-world problems in synthetic biology can be framed as classification tasks, in which combinations of inputs, such as ligand concentrations, pH, or temperature, are mapped to discrete cellular outputs, ranging from reporter fluorescence in whole-cell biosensors [22] to the activation of apoptosis or cytokine release in advanced cell therapies [23, 24]. From a designer perspective, classification can be visualized geometrically by plotting the inputs in a coordinate space (*x*_1_ and *x*_2_ in Fig. 1-A), where the goal is to identify a boundary that separates points belonging to different classes, such as the gray region in Fig. 1-A. Due to their discrete nature, digital circuits generate axis-aligned boundaries (i.e., vertical or horizontal lines in the *x*_1_ − *x*_2_ plane), meaning that generating nonlinear decision boundaries requires layering multiple logic gates, which has been demonstrated in some of the largest and deepest single-cell circuits built to date, including the the 6-input Boolean logic lookup table from [14], and the 12-input ribocomputing device from [25]. Nevertheless, distinguishing between complex biological phenotypes, such as healthy and cancerous cells, typically requires smoother, highly nonlinear decision boundaries [26]. Achieving such boundaries with digital logic would require even more layers of gates, causing the number of genetic components to increase rapidly and limiting the scalability of these circuits for classification tasks.

**Figure 1.**
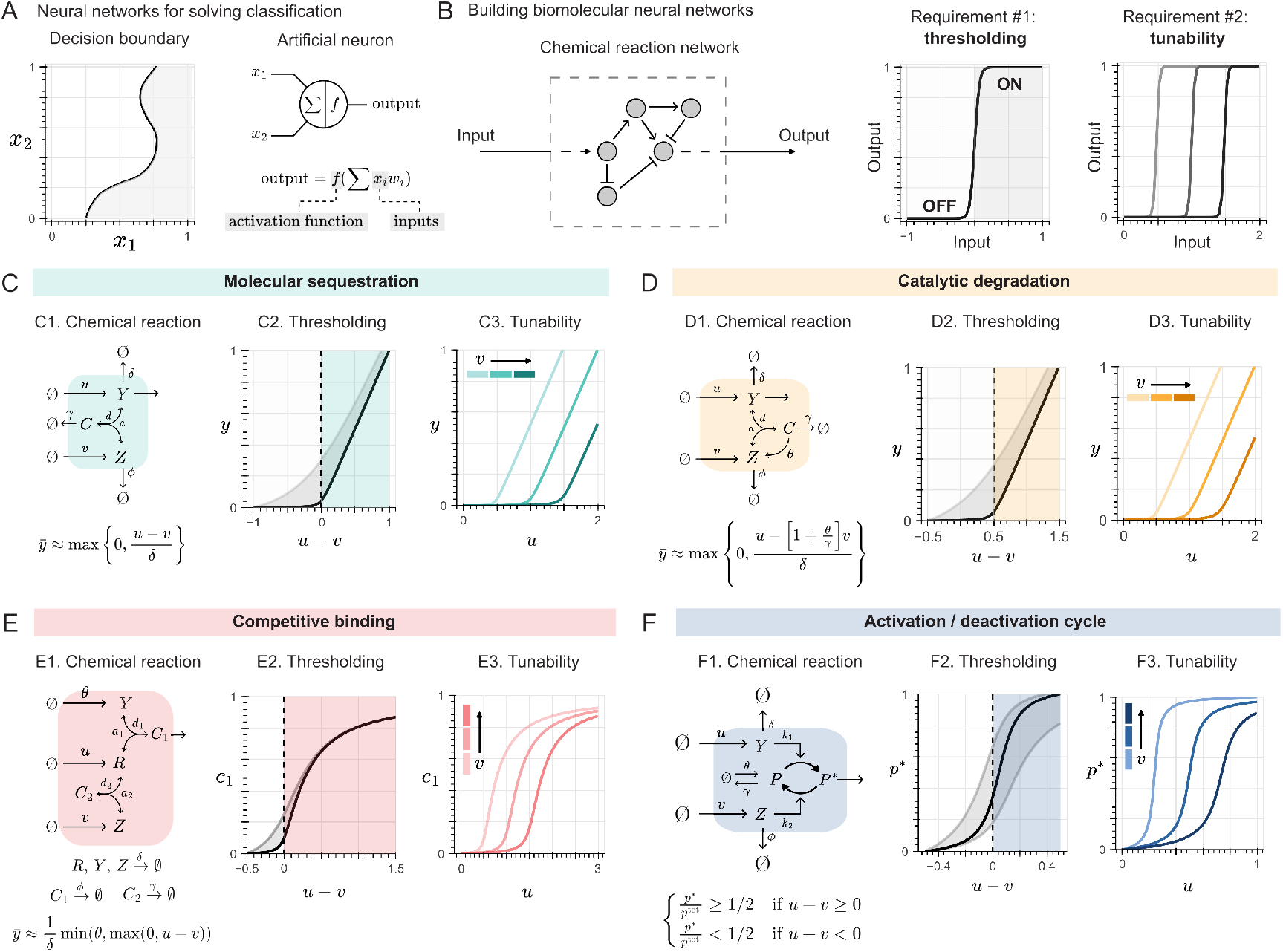
Tunable thresholding as the design principle for building biomolecular neural networks. For classification tasks, multi-layer perceptrons (MLPs) generate decision boundaries (A, black lines) that partition the input space into distinct regions (e.g., white and grey areas). The fundamental computational unit of an MLP is the perceptron, which computes the weighted sum of its inputs (e.g., ∑ x_i_w_i_) and applies an activation function (e.g., f). Analogously, a chemical reaction network whose input–output relationship exhibits a threshold that can be tuned through an accessible parameter can behave as a perceptron (B). In this work, we identify four such networks: molecular sequestration (C1), catalytic degradation (D1), competitive binding (E1), and activation–deactivation cycles (F1). Under specific kinetic conditions, these networks produce thresholding input–output curves (black lines) resembling a Rectified Linear Unit (ReLU) (C2, D2), a saturated ReLU (E2), or a sigmoidal function (F2). When these kinetic constraints are relaxed (grey regions), the networks still display thresholding behavior. Moreover, the thresholds are tunable by adjusting a single rate constant v that controls the production of a sequestering species (C3, E3) or an enzyme (D3, F3).

Due to their expressiveness, neural networks provide a powerful alternative for implementing complex classification tasks in synthetic biology: with sufficient width or depth, they can approximate any continuous function on a bounded domain to arbitrarily small error [27, 28]. Consequently, any decision boundary can, in principle, be realized by an appropriate neural architecture. A few neural network architectures have already been implemented using genetic circuits, and recent studies on the combinatorial logic of cell states suggest that decision boundaries may serve as a natural representation of cellular decision-making [29]. In vitro implementations are based in DNA strand displacement reactions, and include winner (and looser)-takes-all networks, with modules for input weighting, summation, comparison, and signal recovery [30, 31, 32], as well as feedforward [33] and convolutional networks, in which a kernel performs weighted multiplication, with outputs subsequently processed by a feedforward neural network [34, 35]. By contrast, there are only two in vivo implementations to date: a protein-level winner-take-all network using a protease-based architecture [36], and a CRISPR endoribonuclease-based perceptron [37, 38]. Together, the diversity of these implementations highlights the potential for designing next-generation molecular classifiers, with one proof-of-concept network demonstrating the ability to control cell death [36], and raises the question of which principles enable a genetic circuit to function as a neural network.

To explore these principles, we focus on multi-layer perceptrons (MLPs), which are composed of fully connected layers of individual perceptrons, and which we introduced with the name of biomolecular neural networks (BNNs). Each perceptron implements a linear decision boundary, and stacking layers produces nonlinear boundaries. In this work, we examine how candidate chemical reaction networks can implement perceptrons and how these computational motifs can be realized in cells using genetic circuits. We present a quantitative framework that identifies the key feature that enables a chemical reaction network to function as a perceptron: an input–output mapping with a tunable threshold. Using this framework, we derive design rules that show how to construct a suitable decision boundary for a given pattern in the input space (i.e., all the possible input combinations). Ultimately, as a proof-of-principle, we apply the framework *in silico* to design classifiers capable of discriminating healthy and cancer cells based on gene expression data.

## 2 Results

### 2.1 Tunable thresholding as the design principle for biomolecular neural networks

Each perceptron within an MLP operates by feeding the weighted sum of the inputs (∑*x*_*i*_*w*_*i*_ in Fig. 1-A) into a nonlinear function called the *activation function*. For a mathematical function to be consider as such and be useful in a practical setting, so a neural network can be implemented using a reasonable amount of computational resources (i.e., with a fixed number of layers and nodes [39]), it has to be nonlinear (particularly, nonaffine [40]) and exhibit a threshold [41, 28, 27]. These features ensure selective activation by each perceptron: only those units whose inputs exceed their thresholds contribute a nonzero value to the network’s output. However, a fixed threshold constrains the set of functions that the MLP could approximate, also know as its *expressivity*. Hence, to achieve an arbitrarily complex decision boundary, the threshold of the activation function should be tunable by an accessible parameter.

Throughout this work, we assume that a cell’s phenotype can be described through a mathematical function that maps molecular inputs to a discrete cellular outcome, such as proliferation, apoptosis, or differentiation; and to which we refer as *biological function*, following previous terminology [42, 43, 44]. Then, under the universal approximation theorem [27], biomolecular neural networks, could theoretically replicate any biological function synthetically. In the context of classification, depending on whether a combination of inputs lies inside or outside the decision boundary (gray region in Fig. 1-A), a specific phenotypic output is triggered. Then, to build these networks, the input-output relationship of a given chemical reaction network must both exhibit thresholding (Fig. 1-B, left) and allow tuning of that threshold through a downstream mechanism (Fig. 1-B, right), so it can functionally behave as an ideal perceptron.

Here, we identified four chemical reaction networks that are ubiquitous to signaling pathways and gene regulatory networks whose input-output map satisfy the tunable threshold requirement. We generalize each network by assuming different degradation rates for each chemical species to account for the difference in half-lives between DNA, RNA and proteins, as well as homodimers, heterodimers, and molecules after a post-translational modification. We don’t specify the source or number of the inputs nor their specific production rates (*x*_*i*_ and *w*_*i*_, respectively in Fig. 1-A), and consider a “global” production rate and for each chemical species that account for those variables. Then, ∑ *x*_*i*_*w*_*i*_ = *u* and ∑ *x*_*j*_*w*_*j*_ = *v*; in other words, the weighted sum of the inputs *x*_*i*_ and *x*_*j*_, respectively.

The first chemical reaction we identified is molecular sequestration, where two molecules, named *Y* and *Z*, reversibly bind to form an inactive complex (Fig. 1-C1). Examples of molecular sequestration include bacterial toxin-antitoxin systems responsible for antibiotic resistance [45, 46], and the stoichiometric inhibition of transcription factors such as helix-loop-helix (bHLH) and sigma factors [47]. From the mathematical equation describing the network’s output at steady state (Fig. 1-C1), we note that inputs producing the *Y* species are associated with a positive sign, while those producing the *Z* species are associated with a negative sign. The threshold can be identified when mapping the difference between these inputs to the output (Fig. 1-C2); specifically, when the production rates of both species are equal (i.e., *u* − *v* = 0). By considering the steady state concentration of the *Y* species as the output, the input-output map of this chemical reaction network resembles a Rectified Linear Unit (ReLU) activation function when the sequestration rate is fast (or equivalently, when the dissociation constant *K* = *d/a* is very low, such that *K* → 0), as shown by the black line in Fig. 1-C2 [48, 37]. When the sequestration rate is slower due to low-affinity interactions [49, 50, 51, 52], the measured input-output relationship can fall within the gray area in Fig. 1-C2, resembling more a Softplus activation function [53], where the threshold transition is smoother. While the latter has been identified in native regulatory networks [45, 47], the ReLU activation function has been successfully implemented both in vitro and in vivo using synthetic circuits [37, 38]. Regardless of the sequestration regimen, adjusting the production rate *v* of the *Z* species tunes the threshold (Fig. 1-C3), so molecular sequestration satisfies the requirements for an ideal perceptron.

Another input-output response resembling the Softplus activation function has been reported in studies involving miRNA regulation [54, 55, 56, 57] and protease-based synthetic circuits [58, 3], suggesting another candidate for an ideal perceptron. As the second chemical reaction we identify, catalytic degradation of a substrate *Y* by an enzyme *Z* (e.g., the RISC complex recruited by a miRNA, or a protease), with the enzyme recycled back into the reaction (Fig. 1-D1), produces an activation function similar to that of molecular sequestration at steady state (compare Fig. 1-C1, and Fig. 1-D1). Specifically, when the enzyme-substrate binding occurs rapidly, the system approaches a ReLU-like response (black line in Fig. 1-D2) [59], while slower binding kinetics result in a smoother Softplus-like transition (gray area in Fig. 1-D2). As in molecular sequestration, inputs that produce the substrate *Y* are associated with a positive sign, while those that produce the enzyme *Z* are associated with a negative sign. The threshold can be identified when the production rate of the substrate equals a scaled version of the enzyme’s production rate, with the scaling factor determined by the catalytic efficiency of the enzyme. By adjusting the production rate *v* of the enzyme *Z*, the threshold can be tuned (Fig. 1-D3), meaning that catalytic degradation also satisfies the requirements for an ideal perceptron.

Given recent theoretical work on how ligand–receptor interactions enable cell type–specific responses [60], along with how promiscuous protein dimerization enables small networks of monomeric proteins to encode diverse biological functions in homo- and heterodimeric outputs [61], and how a limited set of cytokines and receptors coordinate pathogen recognition during infection [62, 63], the underlying motif governing these interactions emerges as a promising candidate for an ideal perceptron. This mechanism, known as competitive binding, involves a molecule interacting with multiple species at varying affinities, which can be described as “many-to-many” interactions [64]. As the third chemical reaction, we propose a simplified model of competition between two species (*Y* and *Z* in Fig. 1-E1) for a shared molecule (*R* in Fig. 1-E1). The output of interest is the complex *C*_1_, which models, for example, a ligand–receptor complex or an homo-/heterodimer responsible for downstream signaling [60, 61]. The input-output map of this networks, shown in Fig. 1-E2, exhibits two regimes: one with a threshold that occurs when the production rate of one competing species equals that of the shared molecule (i.e., *u* − *v* = 0), and a saturating regime governed by the production of the other competing species (e.g., the production of *Y* at rate *θ* in Fig. 1-E1). For the threshold regime, the inputs producing species *R* are associated with a positive sign, whereas those producing species *Z* are associated with a negative sign. Then, by enforcing non-saturating conditions and for a slower formation of the *C*_1_ complex with respect to the *C*_2_ one (i.e., for *K*_2_ = *d*_2_*/a*_2_ ≪ *K*_1_ = *d*_1_*/a*_1_), this competitive binding motif exhibits an input-output function resembling a ReLU activation function (Fig. 1-E2). This function relaxes into a Softplus shape when the formation of *C*_2_ is slower, shown as the grey area in Fig. 1-E2. Moreover, by adjusting the production rate of the competing species *Z*, the threshold can be tuned (Fig. 1-E3), so this motif satisfies the requirements for an ideal perceptron.

Ultimately, the fourth chemical reaction we identified is generalized as the activation/deactivation cycle (Fig. 1-F1), often used to describe the dynamics between kinase and phosphatase in covalent modification cycles [65], but includes any reversible addition of a chemical group to a molecule [66]. When mapping the inputs to the concentration of the activated protein (*P* ^∗^ in Fig. 1-F1), the sigmoidal curve is obtained (Fig. 1-F2), given a moderate to high binding affinity of each enzyme to its substrate protein (c.f. [67]). For this regime, the threshold corresponds to the input concentration at which the activated protein *P* ^∗^ concentration is more than half of the total protein concentration in the reaction (i.e., *p*^tot^ in Fig. 1-F, left). Moreover, depending on the affinity between the inactivated (or activated) protein and the enzyme that catalyzes its activation (or inactivation, respectively), the shape of the inputoutput curve could vary within the gray area shown in Fig. 1-E2, which is consistent with experimental observations [65, 68, 69]. Regardless of the affinity, the threshold can be observed when the production rate of both species *Y* and *Z* are equal (i.e., *u* − *v* = 0). The inputs that produce the enzyme that activates the protein (i.e., *Y*) are associated with a positive sign, whereas the inputs that produce the enzyme that deactivates the protein (i.e., *Z*) are associated with a negative sign. In addition, by adjusting the production rate *v* of the enzyme *Z* that deactivates the target protein, the threshold can be tuned (Fig. 1-F3), and so the activation/deactivation cycle satisfies the requirement for an ideal perceptron.

### 2.2 Building a multi-layer perceptron

For simplicity, we are going to use the textbook abstraction of a perceptron to represent the chemical reactions detailed in Section 2.1, color coded as shown in Fig. 2-A. To reconstitute the decision boundary in the input space, we screen over increasing values of the network inputs and evaluate the output for each combination, analogous to a titration experiment in biochemistry. Fig. 2-B1 illustrates the linear decision boundary for the sequestration-based perceptron with two inputs, *X*_1_ and *X*_2_, produced at rate constants *w*_1_ and *w*_2_, respectively. Following the notation in Fig. 1-B, we define *u* = *x*_1_*w*_1_ and *v* = *x*_2_*w*_2_. In this framework, the weights of the perceptron are implemented as production rates. Therefore, adjusting them through transcriptional or translational regulatory elements serves as a form of weight tuning that enables control over the slope of the decision boundary. To increase the slope, we could either increase *w*_1_ or decrease *w*_2_, as now a higher concentration of *X*_2_ will be needed to match that of *X*_1_ (Fig. 2-B2). On the other hand, to decrease the slope, we could either decrease *w*_1_ or increase *w*_2_, since now a lesser concentration of *X*_2_ will be needed to match that of *X*_1_ (Fig. 2-B3).

**Figure 2.**
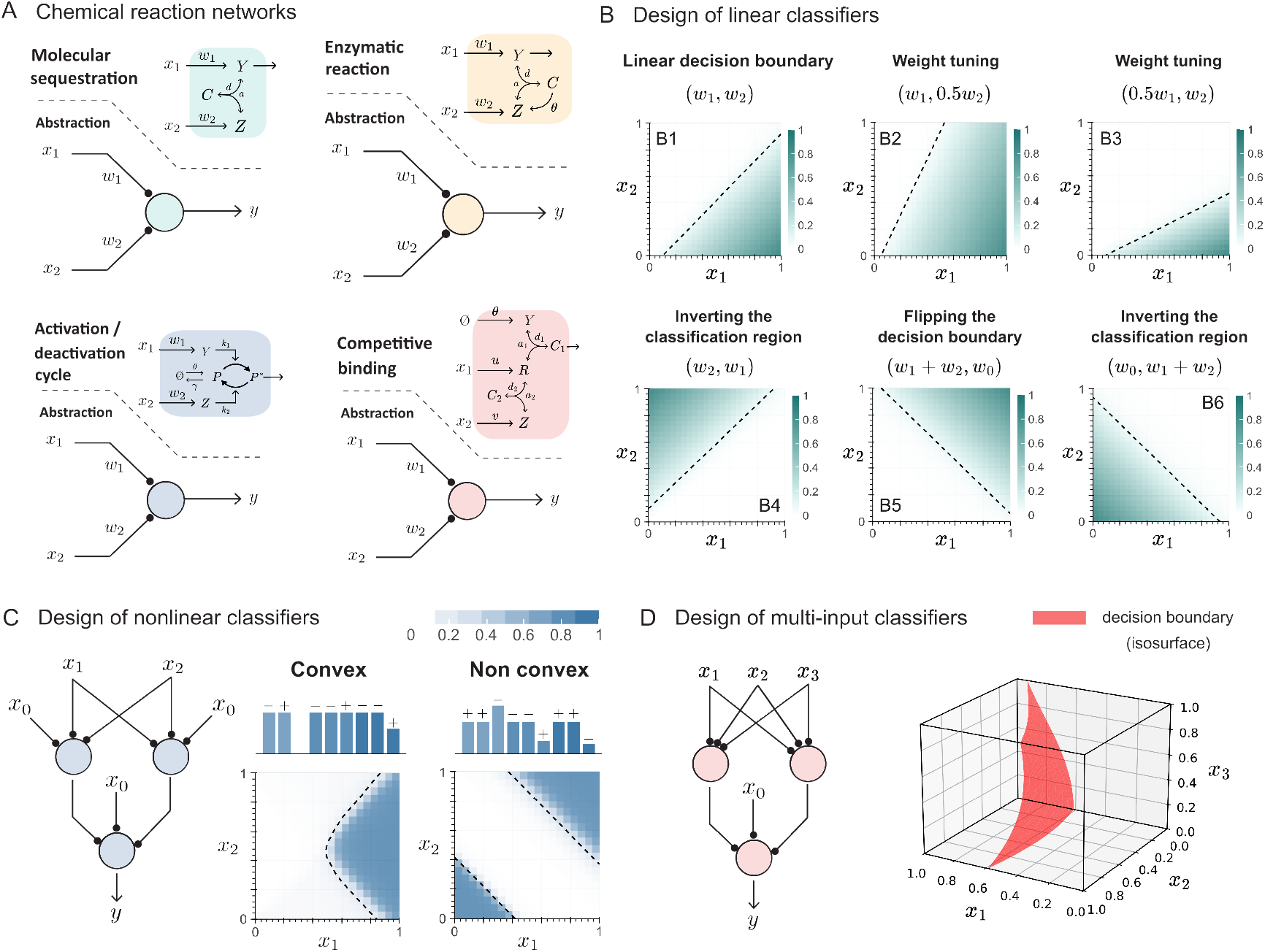
Building multi-layer perceptrons (MLPs) (A) To align with common notation in the machine learning field, we use the symbol of a neuron to summarize the chemical reactions shown to function as ideal perceptrons. Molecular sequestration-based linear classifiers were designed to illustrate how decision boundaries (dashed black lines) define classification regions (green). In Panel B1, the weights were set to w_1_ = 1/h and w_2_ = 1/h, and written above each heatmap as a tuple (w_1_, w_2_). In this notation, the first weight is assigned to a positive effect (i.e., production of the Y species) and the second to a negative effect (i.e., production of the Z species), as determined by their asymptotic approximation. Panels B2–B6 use these weights as the baseline. When two weights are expressed as a sum (e.g., w_1_ +w_2_), it indicates that two inputs act in combination to produce a species within the chemical reaction network. (C) MLPs based on the activation/deactivation network were designed, illustrating nonlinear decision boundary (black dashed lines) that define a classification region (blue) that can be either convex or non-convex. An architecture with two nodes in the hidden layer and a single node in the output layer is considered. The bar heights above each heatmap indicate the magnitude of the weights associated with each chemical species, and the signs represent if the inputs produce either the species Y (i.e., a positive sign) or the species Z (i.e., a negative sign). (D) An MLP based on the competitive binding network was designed to illustrate a nonlinear decision boundary in a 3D input space (i.e., when considering three inputs x_1_, x_2_ and x_3_). For both (B) and (C), heatmap values were normalized by dividing each value by the maximum within each simulation. A grid size of N = 20 was used for both inputs x_1_ and x_2_, with concentrations ranging from 0 to 1 µM. For (D), the decision boundary was estimated by computing the isosurface at the threshold value determine by the threshold in the output node.

For every chemical reaction, the sign of the weights are encoded in the species that each input produces (Fig. 1-C to F). For example, in the sequestration-based perceptron shown in Fig. 2-A, the reaction 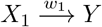 associates *w*_1_ with a positive sign, whereas the reaction 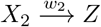 associates *w*_2_ with a negative sign. Inverting the region within the input space where the perceptron’s output is nonzero (i.e., the *classification region*) requires switching the species that each input produces. For the same example as before, switching the inputs so that *X*_1_ produces species *Z* at rate constant *w*_1_ and *X*_2_ produces species *Y* at rate constant *w*_2_ yields the decision boundary shown in Fig. 2-B4. Moreover, considering an additional constant input *X*_0_, also known as the *bias*, can extend or decrease the classification region, depending on the species it produces. More importantly, the addition of this species enables flipping the decision boundary by assigning positive weights to *x*_1_ and *x*_2_, and a negative weight to *x*_0_, which is produced at rate constant *w*_0_. This configuration results in the decision boundary shown in Fig. 2-B5. Similar to the switching shown in Fig. 2-B4, assigning the positive weight to *x*_0_ and negative weights to both *x*_1_ and *x*_2_ enables the inversion of the classification region, as shown in Fig. 2-B6. A similar analysis for the rest of the chemical reactions shown in Fig. 2-B is available in the Supplementary Figure 1.

Furthermore, by layering individual perceptrons into an MLP architecture, we can construct nonlinear decision boundaries. Fig. 2-C illustrates an MLP based on the activation/deactivation cycle, with two inputs (*X*_1_ and *X*_2_), a *hidden layer* of two nodes (each with a bias species *X*_0_), and a single output node with its own bias. This same architecture can yield different decision boundaries by adjusting the weights and choosing which species each input produces, following the principles outlined in Fig. 2-B. In Fig. 2-C, we show two examples: a convex boundary (left) and a non-convex one (right). Above each heatmap, schematics depict the relative weight magnitudes (via bar height) and signs (positive for producing the activating species *Y*, and negative for the inactivating species *Z*). Additionally, since the network is small, with a total of 3 nodes, the resulting boundaries remain interpretable: we can directly map the output to the linear regions defined by the hidden nodes. For example, for the non convex case, the output of the nodes in the hidden layer are those shown in Fig. 2-B5 and B6, which are summed in the output layer to produce the nonlinear decision boundary capable of solving the XOR problem [70]. A larger repertoire of nonlinear decision boundaries based on the architecture illustrated in Fig. 2-C are available in Supplementary Figures 2 to 4.

So far, we have used a two-dimensional input space to design decision boundaries, as it provides the minimal number of inputs needed to generate the chemical species required for thresholding behavior. However, biomolecular neural networks can operate with more than two inputs. These inputs do not need to be strictly chemical in nature; for example, they could represent mechanical signals, light, or other types of stimuli [71, 72]. The design of such multi-input classifiers follows the same principles described earlier. Figure 2-D (left) shows an MLP based on the competitive binding mechanism that receives three inputs (*X*_1_, *X*_2_, and *X*_3_), which are processed through a hidden layer with two nodes. The outputs of these nodes, together with a bias species, are then passed to the output layer to generate the decision boundary shown in Fig. 2-D (right). The ability to translate, rotate, and flip the decision boundary by tuning the weights is illustrated in Supplementary Figure 5.

### 2.3 Rational design of biomolecular neural networks for cancer cell classification

As a proof of principle, here we focus on strategies based on biomolecular neural networks to augment the classification capabilities of cell-based cancer therapies, as an illustrative example of cell state sensing. Our analysis is not intended to be comprehensive, but rather serve as an exploratory effort to identify patterns among biomarkers and examine the utility and limitations of the proposed biomolecular neural networks. We used the publicly available dataset provided by Dannenfelser et al (2020) [73], which consists of batch-corrected and normalized gene expression data assembled from the Cancer Genome Atlas (TCGA) and the Genotype-Tissue Expression (GTEx) project. The dataset corresponds to 33 cancer types and 30 healthy tissues. As previously annotated, tissue-specific comparisons for each cancer type were not possible due to the lack of a direct mapping between each cancer type and its tissue of origin. To address this, we screened the TCGA dataset to identify the tissue from which each cancer sample was derived and manually annotated the corresponding tissue type (see Supplementary Table 1). For example, expression data from lung adenocarcinoma and lung squamous cell carcinoma were grouped under the “lung cancer” category.

After this annotation step, we analyzed the expression levels of each gene in a pair (each referred to as an antigen) across tissue types, creating two-dimensional visualizations of their distributions, which we call *patterns*. We only analyzed 19 out of the 30 available tissue types after filtering by sample size and class balance (Supplementary Figure 6). In principle, given a certain pattern, biomolecular neural networks can approximate an adequate decision boundary, given the sufficient number of nodes in the hidden layer (i.e., it’s width). We aim not only to design a classifier with the highest performance, but the smallest circuit as well, driven by experimental feasibility. Therefore, we took an incremental approach, starting with linear classifiers, and then moving into nonlinear classifiers with 2 nodes in the hidden layers, then 3 nodes, and finally 4 nodes. The rationale for stopping at this width is addressed in the Discussion. For the linear classifiers, we used the activation/deactivation reaction given that it maps the linear combination of the inputs into a value that can be interpreted as a probability through a sigmoid-like function (see Figure 1-F). For the nonlinear classifiers, we used sequestration-based perceptrons in the hidden layer, and the activation/deactivation reaction for the output layer.

Given that designing one classifier per pattern will be computationally expensive, we adopted a heuristic approach to guide our exploration of the antigen space to find representative examples from the dataset that could highlight the advantages and limitations of the proposed networks. We computed a separability metric for each antigen, under the assumption that antigens with higher values would, when combined in pairs, require simpler decision boundaries to achieve high performance [74, 75, 76, 77], provided that a single antigen is often not enough to separate between healthy and cancer cells (see exploratory analysis in Supplementary Figure 7). Therefore, we computed the Earth’s Mover Distance (EMD, c.f. [74]) per antigen, and sorted the list of antigens per tissue type. Finally, we divided the EMD distribution across antigens into three groups (high, medium and low EMD), and evaluated patterns that emerged from antigen combinations within the top 10 candidates in the same group, and between groups (see Supplementary Figure 8). Ultimately, we constructed confusion matrices and evaluated performance using precision, recall, F1 score, and the area under the ROC curve (AUC).

We primarily report the F1 score, which balances precision and recall and is therefore appropriate for our dataset (Supplementary Figure 6). Across all tissues, combinations containing at least one high-EMD antigen achieved a median F1 score of 0.939 (IQR 0.091), compared to 0.826 (IQR 0.260) for pairs without any high-EMD antigen. A Wilcoxon signed-rank test confirmed that F1 scores were significantly higher for high-EMD combinations than for non-high-EMD combinations across tissues (*p <* 0.001). Figure 3-A illustrates an example from lung tissue using SCNN1B and EDNRB, which have been previously identified as biomarkers for lung adenocarcinoma [78, 79, 80]. Although high-EMD antigens often exhibit higher performance (Supplementary Figure 9), they are not a guarantee. We identified outliers with F1 scores falling below 0.3, corresponding to antigen pairs with substantial overlap between healthy and cancer samples when considered jointly. Medium- and low-EMD combinations showed similar overlap, indicating that linear classifiers are insufficient to separate them and that an architecture capable of generating nonlinear decision boundaries is required. For these patterns, increasing the network complexity from a single perceptron to an MLP with 2 nodes in the hidden layer significantly improved F1 scores. Across all tissues, the difference in F1 score between this MLP architecture and the individual perceptron (denoted by the metric Δ*F*_1_) was consistently positive (mean Δ*F*_1_ = 0.049, median Δ*F*_1_ = 0.017), and a Wilcoxon signed-rank test confirmed that this improvement was statistically significant (*p <* 0.001. See also Supplementary Table 2 and Supplementary Figure 10). Figure 3-B shows an illustrative example from blood tissue using the antigens ANPEP and PVRL1, which correspond to an outlier of the High-Medium category. In this case, the linear classifier is unable to separate the two classes (F1 ≈ 0), whereas the MLP with 2 nodes in the hidden layer successfully captures the nonlinearity in the pattern, achieving an F1= 0.781.

**Figure 3.**
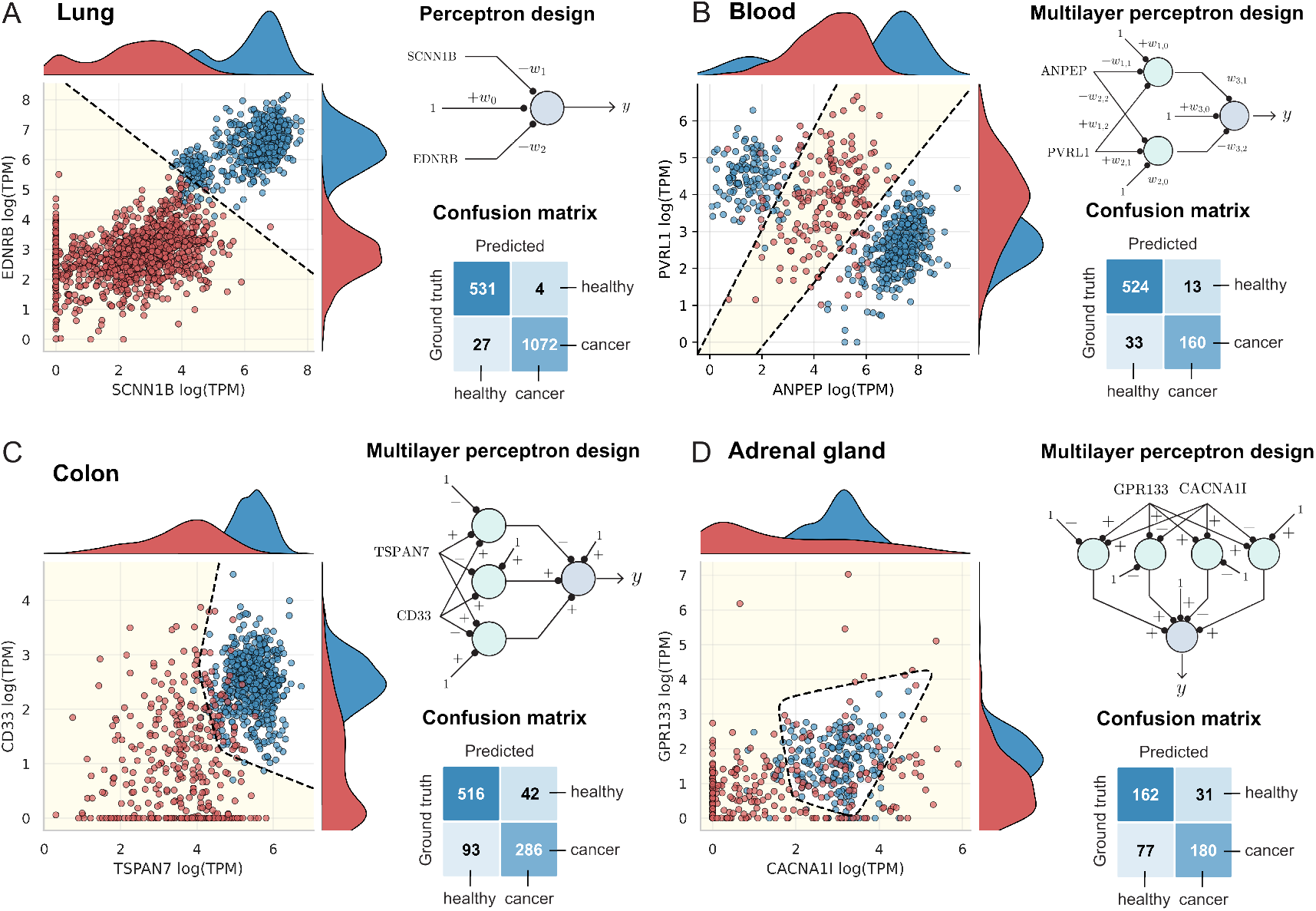
Design of tissue-specific linear and nonlinear classifiers for antigen recognition in cancer therapeutics. The linear decision boundary of each sequestration-based classifier is shown as a dashed black line, with the region classified as cancer highlighted in yellow. The true labels of the samples are color-coded, with cancer samples in red and healthy samples in blue. The sign of each weight (+ or −) was determined from either the activation/deactivation cycle’s (blue) or the molecular sequestration’s (green) asymptotic approximation (see Figure 1-C and F, respectively).

When increasing the number of nodes in the hidden layer, we observed a small but consistent increase in F1 score across tissues (2-to-3 nodes: median Δ*F*_1_ = 0.0042 for Medium-Medium pairs, Wilcoxon signed-rank test *p <* 0.001; 3-to-4 nodes: median Δ*F*_1_ = 0.0022, *p <* 0.001). While most nonlinear patterns in our dataset were captured using an MLP with 2 nodes in the hidden layer (Supplementary Table 2), some antigen combinations required wider networks, especially those with a Medium-EMD antigens (Supplementary Figures 16–20). Fig. 3-C shows colon tissue using the TSPAN7 and CD33 antigens from the Medium-Medium category. Using an MLP with 3 nodes in the hidden layer, we captured the pattern with F1 = 0.9004. Similarly, Fig. 3-D shows adrenal gland using the CACNA1I and GPR133 antigens from the Medium-Low category. Using an MLP with 4 nodes in the hidden layer, we captured the pattern with F1 = 0.7505. More examples are available in Supplementary Figure 21. Nevertheless, for Low-Low antigen combinations, even the most complex neural networks designed (i.e., 4 nodes in the hidden layer) had limited performance compared to other antigen combinations: the median F1 across tissues was 0.756, and the 90th percentile was 0.88. In contrast, the 90th percentile for Medium-Medium, Medium-Low, and High-containing pairs ranged from 0.92 to 0.95, highlighting that Low-Low combinations remain the most difficult patterns to separate in our dataset. Addressing this challenge is left for future work.

## 3 Discussion

Cells have evolved to sense a wide range of input combinations and integrate those signals through signaling pathways to produce context-specific responses, such as differentiation, cell-type specification, and patterning [81, 82]. To replicate this information-processing capacity, synthetic biology has developed large-scale circuitry inspired by the fundamental principles of computer science. Within this framework, neuromorphic computing implemented using genetic circuits offers the opportunity to significantly enhance the computational capabilities of single cells. Here, we focused on formalizing the design principles that enable a candidate genetic circuit to operate as a perceptron, the fundamental unit of a multi-layer perceptron, which constitutes the core implementation of modern neural network architectures. We further identified four chemical reaction networks that comprise the majority of circuits available in the current repertoire of synthetic biology parts: molecular sequestration, already proven to operate as a perceptron [37, 38]; catalytic degradation reactions, well-studied in the context of miRNAs [83] and protease-based circuits [58]; activation/deactivation cycles, implemented as phosphorylation/dephosphorylation reactions for feedback control [68, 69]; and competitive binding reactions, which, although explored in some synthetic biology implementations, are more commonly observed in natural signaling pathways [61]. This observation raises the intriguing question of whether neuromorphic computing could be extended to native signaling pathways and gene regulatory networks [29]. Overall, these genetic circuits could be repurposed using available genetic engineering tools to operate as perceptrons, following the tunable thresholding principle outlined in this work.

In the context of classification, differentiating between phenotypes involves identifying a set of genes that uniquely define one of the states of interest, and the difficulty of this task can vary substantially. Some phenotype distinctions could be achieved using a small subset of biomarkers through cell-type specific enhancers [84], or characteristic miRNA expression profiles [83]. However, the vast majority of cases are considerably more challenging. For example, distinguishing healthy from cancerous cells originating from the same tissue is difficult because their expression profiles highly overlap (c.f. [85], observed in Fig. 3 and Supplementary Figures 13-15). At higher levels of complexity, classification requires mining high-dimensional datasets to identify a minimal yet informative set of biomarkers [73], with only rare situations where one marker is sufficient. Since genetic circuits process a finite number of inputs, dimensionality must be reduced to an even smaller set of biomarkers, which introduces nonlinearity to the classification task. Biomolecular neural networks are particularly useful in this setting because they can be engineered to match the nonlinear decision boundaries present in the data, eliminating the need to reshape the dataset to fit the limited expressive capacity of non-neuromorphic genetic circuit architectures.

From a practical perspective, the experimental implementation of biomolecular neural networks reveals a challenge regarding their predicted expressivity: there is an upper limit to the number of perceptrons that can be stably delivered into a cell using current DNA delivery methods, which is even more restrictive for primary cell lines intended for therapeutic applications [86]. In other words, the limited number of perceptrons restricts the shapes of decision boundaries that can be reconstructed. Interestingly, a similar constraint exists in native signaling pathways, where the finite size of the genome limits the number of genes available for computation, yet cells exhibit remarkable computational capabilities [29]. Beyond engineering more compact genetic circuitry, future work could take inspiration from strategies that cells use to increase computational capacity without expanding the number of genetic elements, in contrast to artificial neural networks where higher expressivity is typically achieved by increasing network size. Such strategies may include many-to-many protein interactions [61, 64], in which heterogeneous binding affinities might be acting as intrinsic hidden layers; stochasticity, acting as a regularizer [48], and nonlinearities arising from competition over shared cellular resources [87]. Understanding and leveraging these mechanisms could inform the design of next-generation biomolecular neural networks, though further research is needed. Furthermore, we didn’t address the online training of these networks as it’s left for future work, yet we acknowledge current advances towards a chemical learning framework [88].

Ultimately, the tunable thresholding principle holds mathematically and can be used to explain the mechanisms behind existing neural network implementations (including DNA strand displacement reactions [30, 31, 32, 33, 35], captured by the molecular sequestration model; as well as convolutional neural networks that rely on enzymes [34] and protease-based circuits [36], both captured by the catalytic degradation model), several considerations should be taken into account regarding the scope of the *in silico* cancer cell classifiers. First, the classifiers rely on a subset of the TCGA and GTEx harmonized datasets, which introduces class imbalance and, in some cases, artificially skewed gene expression distributions (see Supplementary Information, Section 4). Second, the input units used for offline training were transcripts per million, which do not directly correspond to protein concentrations and require assumptions about total mRNA per cell and uniform capture efficiency to estimate absolute transcript levels [89]. Consequently, the decision boundaries reconstructed here should be interpreted as a proof of principle for applying biomolecular neural networks to cancer cell classification, rather than as a method for antigen discovery. Ultimately, we did not account for the biological nature of the inputs used to design the biomolecular neural networks. Some inputs correspond to receptors with extracellular domains, but we did not verify whether these or other candidate biomarkers could be sensed using currently available engineered receptors or related technologies. Nevertheless, we anticipate that advances in protein engineering could enable the detection of these molecules if they prove to be valuable biomarkers for cancer classification [90]. In practice, incorporating such a sensing module would effectively add an input layer to the network, which needs to be considered for circuit design.

