## Supplementary Information for "Design principles of neuromorphic computing using genetic circuits"

### Design principles of chemical-based neuromorphic systems

December 1, 2025

### 1 Chemical reactions and dynamical models

#### 1.1 Molecular sequestration

The molecular sequestration reaction involves two chemical species $Y$ and $Z$, produced at rates $u$ and $v$, respectively, that interact by reversibly binding at rate $a$ to form an inactive complex $C$ that dissociates at rate $d$. This interaction can be modeled with the following chemical reactions:

$$
\begin{array}{lll}
\emptyset \xrightarrow{u} Y & \emptyset \xrightarrow{v} Z & \text{Production} \\
Y + Z \underset{d}{\overset{a}{\rightleftharpoons}} C & & \text{Sequestration (binding/unbinding)} \\
Y \xrightarrow{\delta} \emptyset & & \text{Decay} \\
Z \xrightarrow{\phi} \emptyset & C \xrightarrow{\gamma} \emptyset & \text{Decay}
\end{array} \tag{1}
$$

Using the law of mass action, we can write down the Ordinary Differential Equations (ODEs) that describe the dynamics of the chemical reactions in (1):

$$\dot{y} = u + dc - ayz - \delta y \tag{2}$$
$$\dot{z} = v + dc - ayz - \phi z \tag{3}$$
$$\dot{c} = ayz - dc - \gamma c \tag{4}$$

#### 1.2 Catalytic degradation

We focus on a mechanism in which an enzyme ($Z$) mediates the degradation of its substrate ($Y$). The free enzyme reversibly binds to the substrate at rate $a$, forming an enzyme–substrate complex. This complex can dissociate at rate $d$, while simultaneously facilitating the degradation of the substrate. After dissociation, the enzyme is effectively "recycled" into the reaction pool at rate $\theta$.This interaction can be modeled with the following chemical reactions:

$$
\begin{array}{lll}
\emptyset \xrightarrow{u} Y & \emptyset \xrightarrow{v} Z & \text{Production} \\
Y + Z \underset{d}{\overset{a}{\rightleftharpoons}} C & & \text{Enzyme:substrate complex formation} \\
C \xrightarrow{\theta} Z & C \xrightarrow{\gamma} \emptyset & \text{Recycling / Decay} \\
Z \xrightarrow{\phi} \emptyset & Y \xrightarrow{\delta} \emptyset & \text{Decay}
\end{array} \tag{5}
$$

Using the law of mass action, we can write down the Ordinary Differential Equations (ODEs) that describe the dynamics of these reactions

$$\dot{y} = u - \delta y - ayz + dc \tag{6}$$
$$\dot{z} = v - \phi z - ayz + c(d + \theta) \tag{7}$$
$$\dot{c} = ayz - c(\theta + \gamma + d) \tag{8}$$

#### 1.3 Competitive binding

We model a simplified reaction where two species compete for a "shared" one by considering a chemical species $R$, produced at a rate $u$, that is able to reversibly bind at rate constant $a_1$ with another species $Y$, produced at a rate constant $\theta$, to form a complex $C_1$ that dissociates at rate constant $d_1$. Simultaneously, a chemical species $Z$, produced at rate $v$, is able to reversibly bind with $R$ at rate constant $a_2$ to form another complex $C_2$ that dissociates at rate constant $d_2$. In this scheme, $Y$ and $Z$ compete for $R$. These interactions can be modeled using the following chemical reactions:

$$\begin{array}{lll} \emptyset \xrightarrow{u} R & R \xrightarrow{\delta} \emptyset & \text{Production/Decay} \\ \emptyset \xrightarrow{\theta} Y & Y \xrightarrow{\delta} \emptyset & \text{Production/Decay} \\ \emptyset \xrightarrow{v} Z & Z \xrightarrow{\delta} \emptyset & \text{Production/Decay} \\ R + Y \underset{d_1}{\overset{a_1}{\rightleftharpoons}} C_1 & R + Z \underset{d_2}{\overset{a_2}{\rightleftharpoons}} C_2 & \text{Complex formation} \\ C_1 \xrightarrow{\phi} \emptyset & C_2 \xrightarrow{\gamma} \emptyset & \text{Decay} \end{array} \tag{9}$$

Using the law of mass action, we can write down the ODEs that describe the dynamics of these reactions as follows:

$$\dot{r} = u - \delta r + (d_1 c_1 - a_1 r y) + (d_2 c_2 - a_2 r z) \tag{10}$$
$$\dot{y} = \theta - \delta y + (d_1 c_1 - a_1 r y) \tag{11}$$
$$\dot{z} = v - \delta z + (d_2 c_2 - a_2 r z) \tag{12}$$
$$\dot{c}_1 = (a_1 r y - d_1 c_1) - \phi c_1 \tag{13}$$
$$\dot{c}_2 = (a_2 r z - d_2 c_2) - \gamma c_2 \tag{14}$$

#### 1.4 Activation/deactivation reactions

We model activation and deactivation cycles by considering a protein $P$, produced at a rate $\theta$, that can be activated by a protein $Y$, produced at a rate $u$, and deactivated by a protein $Z$, produced at a rate $v$.

$$\emptyset \xrightarrow{\theta} P\,, \quad \emptyset \xrightarrow{u} Y\,, \quad \emptyset \xrightarrow{v} Z \tag{15}$$

The proteins $Y$ and $P$ can bind at rate constant $a_1$ to form an intermediate complex $C_1$ that dissociates at rate $d_1$. This complex yields the activated protein $P^*$ at rate constant $k_1$

$$Y + P \underset{d_1}{\overset{a_1}{\rightleftharpoons}} C_1 \xrightarrow{k_1} Y + P^* \tag{16}$$

Similarly, the protein $Z$ and the activated protein $P^*$ can bind at a rate constant $a_2$ to form an intermediate complex $C_2$ that dissociates at rate $d_2$. This complex yields back the protein $Y$ (i.e., inactivated) at a rate constant $k_2$

$$Z + P^* \underset{d_2}{\overset{a_2}{\rightleftharpoons}} C_2 \xrightarrow{k_2} Z + P \tag{17}$$

The (inactivated) protein $P$, the proteins $Y$ and $Z$, and both complexes $C_1$ and $C_2$ are subject to degradation at rate $\delta$, whereas the activated protein $\phi$ is consider to have a different degradation $\phi$. For some covalent modification cycles, we note that $\delta = \phi$, whereas for those involving downstream degradation $\delta \neq \phi$.

$$P, Y, Z, C_1, C_2 \xrightarrow{\delta} \emptyset\ , P^* \xrightarrow{\phi} \emptyset \tag{18}$$

Using the law of mass action, we can write down the Ordinary Differential Equations (ODEs) that describe the dynamics of these reactions

$$
\begin{aligned}
\dot{y} &= u - \delta y - a_1 y p + (d_1 + k_1) c_1 && (19) \\
\dot{z} &= v - \delta z - a_2 z p^* + (d_2 + k_2) c_2 && (20) \\
\dot{c}_1 &= a_1 y p - (d_1 + k_1) c_1 - \delta c_1 && (21) \\
\dot{c}_2 &= a_2 z p^* - (d_2 + k_2) c_2 - \delta c_2 && (22) \\
\dot{p} &= \theta - \delta p - a_1 y p + d_1 c_1 + k_2 c_2 && (23) \\
\dot{p}^* &= -\phi p^* - a_2 z p^* + d_2 c_2 + k_1 c_1 && (24)
\end{aligned}
$$

### 2 Design of linear classifiers

Since the input–output response of each chemical reaction network detailed in Section 1 satisfies the requirement of tunable thresholding, we can build an ideal perceptron that operates as a linear classifier using any of these networks. The ability to adjust the decision boundary by tuning the weights, to reverse the classification region by switching the inputs, and to flip the boundary by introducing a constant bias species was shown in Supplementary Figure 2 of the Main Text for the sequestration-based implementation. In Supplementary Figure 1, we show that all of these features can also be achieved with the other networks, with the sequestration-based case included in the first row for comparison.

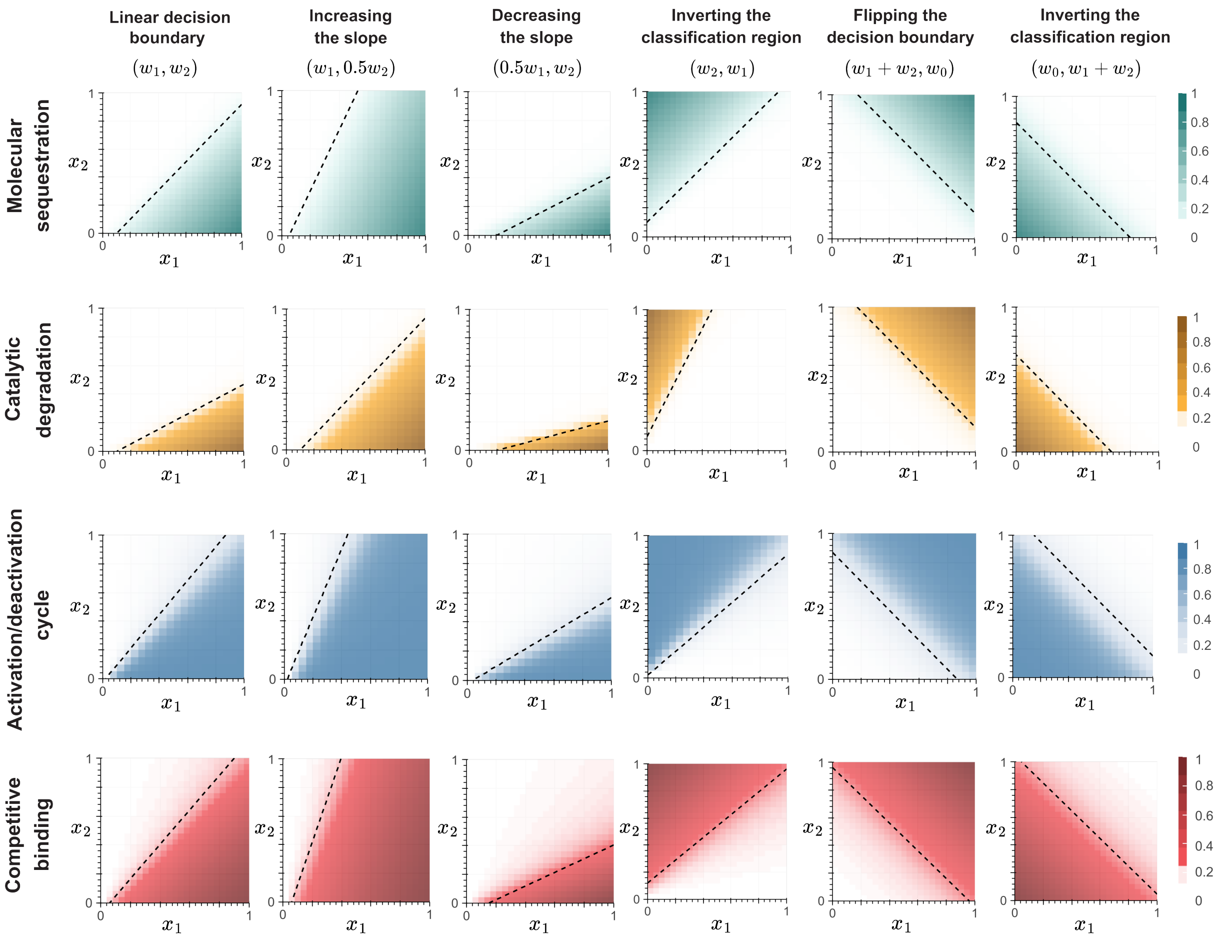

***Figure 1: Design of linear classifiers using chemical reaction networks with tunable thresholds.*** *For all chemical reactions, the weights were set to $w_1 = 1/h$ and $w_2 = 1/h$, and written above each heatmap as a tuple ($w_1$, $w_2$). In this notation, the first weight is assigned to a positive effect (i.e., production of the $Y$ species) and the second to a negative effect (i.e., production of the $Z$ species), as determined by their asymptotic approximation. Simulations from the second column onwards use these weights as the baseline. When two weights are expressed as a sum (e.g., $w_1 + w_2$), it indicates that two inputs act in combination to produce a species within the chemical reaction network. All heatmaps were normalized by dividing each value by the maximum within each simulation. A grid size of $N = 20$ was used for both inputs $x_1$ and $x_2$, with concentrations ranging from 0 to 1 $\mu$M. The decision boundary for each implementation is represented by black dashed lines overlaid on the heatmaps.*

### 3 Design of nonlinear classifiers

Nonlinear decision boundaries are generated by layering individual perceptrons into what is known as a multi-layer perceptron (MLP) architecture. In this work, we focus on fully connected MLPs composed of a single hidden layer and an output layer, where each node (i.e., perceptron) is connected to all nodes in the subsequent layer, but not to other nodes within its own layer. The resulting decision boundaries were further classified into two categories: *convex* (see Supplementary Figure 2), where the classification region is closed (that is, any two points $(x_1, x_2)$ within the region can be connected by a line segment that lies entirely within the region), and *non-convex* (see Supplementary Figure 3), where this condition does not hold. Note that the decision boundaries presented in Figs. 2 and 3 **do not** represent the complete expressive power of an MLP with two hidden nodes and a single output node, but were instead selected as illustrative examples of arbitrary nonlinear patterns.

#### 3.1 Convex decision boundaries

As a first case study, we design an MLP to generate a decision boundary with a polygonal (convex) shape. Supplementary Figure 2-A (left) shows a network composed of two hidden-layer nodes and a single output node, implemented using the catalytic degradation network. As described in the Main Text, the sign of each weight reflects its asymptotic effect in steady state: connections to species $Y$ are assigned a positive sign $(+)$, while those to species $Z$ are assigned a negative sign $(-)$. The magnitude of each weight is shown in the bar plot to the right of the schematic; the absence of a bar indicates that no bias species $X_0$ was included for that node. The corresponding heatmaps show the decision boundaries produced by each node. Nodes 1 and 2 produce linear decision boundaries, which were designed so that their classification regions intersect. When both are connected to the output node with positive weights, the resulting nonlinear decision boundary approximates a scaled sum of their individual classification regions. The size of this intersection is modulated by the weight associated with the bias species producing $Z$ (i.e., the enzyme that catalyzes the degradation of the substrate $Y$).

As a second case study, we consider the same MLP architecture, now implemented using the network based on the activation/deactivation cycle. In this configuration, the inputs to the output node were inverted (see Supplementary Figure 2-B, left), and thus assigned the opposite sign. The weights were also adjusted so that the decision boundaries of Nodes 1 and 2 match those in the previous case. As a result, the output at Node 3 corresponds to the geometric complement of the classification region observed in the first case (compare Node 3 in Supplementary Figure 2-A and B).

In a third case (Supplementary Figure 2-C, left), we extended the previous design by including a bias species in Node 1. This modification was used to generate opposing linear decision boundaries for Nodes 1 and 2. If the outputs of both nodes are connected to the output node with positive weights, the resulting output corresponds to a scaled sum of their classification regions, forming a non-convex decision boundary. Instead, by assigning negative weights to both connections, the network computes the geometric complement of that region, resulting in an angled, band-like decision boundary that spans the input space.

#### 3.2 Non convex decision boundaries

The design of non-convex decision boundaries using the same architecture described in Section 3.1 (i.e., two hidden-layer nodes and a single output node) can be achieved by configuring the hidden-layer nodes to generate linear decision boundaries with opposing classification regions. Two illustrative examples are shown in Supplementary Figure 3. The first architecture, shown in Supplementary Figure 3-A, is the same

MLP architecture proposed in Supplementary Figure 2-C, but with inverted inputs to the output node. As a result, the nonlinear decision boundary corresponds to the geometric complement of the band-like decision boindary, as shown for Node 3 in Supplementary Figure 3. The second architecture, shown in Supplementary Figure 3-B, represents the three-node network with the minimal number of molecular species capable of producing a nonlinear decision boundary.

##### 3.3 Increasing the nodes in the hidden layer

In principle, the expressivity of an MLP increases with the number of nodes in its hidden layer [1, 2]. To illustrate this in the context of biomolecular neural networks, we implemented a sequestration-based MLP with three hidden nodes and one output node, as shown in Supplementary Figure 4 (top-left). Each hidden node was designed so that its decision boundary, shown as the output of Node 4 in Supplementary Figure 4 (bottom-right), forms a closed-loop region that exemplifies the classification of a nonlinear pattern. Moreover, building on the intuition developed in Sections 3.1 and 3.2 for constructing convex and non-convex boundaries, we could invert the classification region within the closed-loop by correspondingly inverting the order of the inputs to Node 4. This corresponds to connecting the outputs of the hidden-layer nodes to the production of the corresponding $Z$ species in Node 4, while the bias species $x_0$ drives the production of the corresponding $Y$ species.

##### 3.4 Decision boundaries for three-input classifiers

Similar to artificial neural networks, biomolecular MLPs can handle multiple inputs as long as the transcriptional unit controlling the genetic circuit can process them. This can be achieved, for example, using multi-input promoters [3, 4, 5] or by leveraging alternative splicing [6]. In Supplementary Figure 5-A, we illustrate a linear decision boundary formed by a single perceptron, shown in red within a 3D input space. In contrast, Supplementary Figure 5-B presents a nonlinear boundary generated by a network architecture with two nodes in the hidden layer and one node in the output.

The same design principles used for 2D classifiers can be extended to 3D by projecting onto each axis. However, scaling this approach to even higher dimensions is challenging, both due to biological constraints on multi-input processing and the absence of suitable learning algorithms instead of rational design. Exploring architectures with multiple outputs remains an open direction for future work.

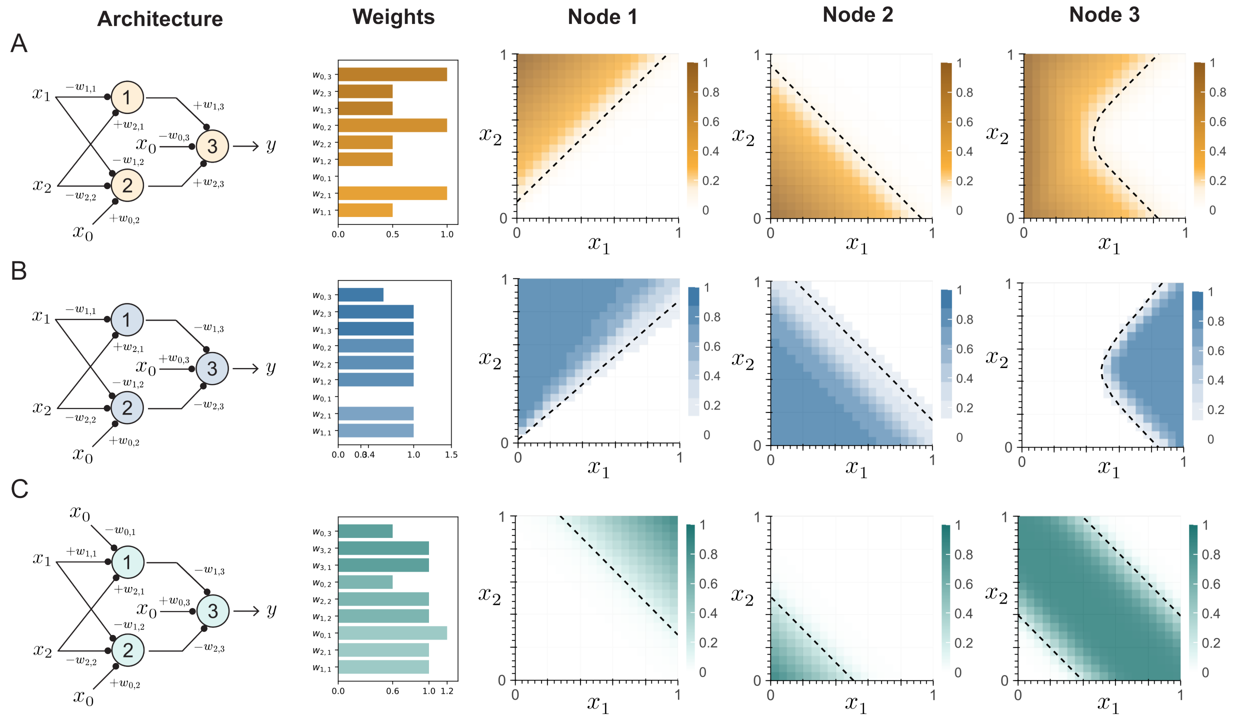

***Figure 2: Design of nonlinear classifiers with a convex decision boundary***. *Multi-layer perceptrons (MLPs) with 2 nodes in the hidden layer and one node as the output layer were designed based on the catalytic degradation (A), activation/deactivation cycle (B) and molecular sequestration reactions (C). The sign of each weight ($+$ or $-$) was determined from its asymptotic approximation, while the corresponding magnitudes are shown in the bar plots. The absence of a bar indicates that no bias species was included. All heatmaps were normalized by dividing each value by the maximum within each simulation. A grid size of $N = 20$ was used for both inputs $x_1$ and $x_2$, with concentrations ranging from 0 to 1 $\mu$M. Decision boundaries for each implementation are shown as black dashed lines overlaid on the heatmaps.*

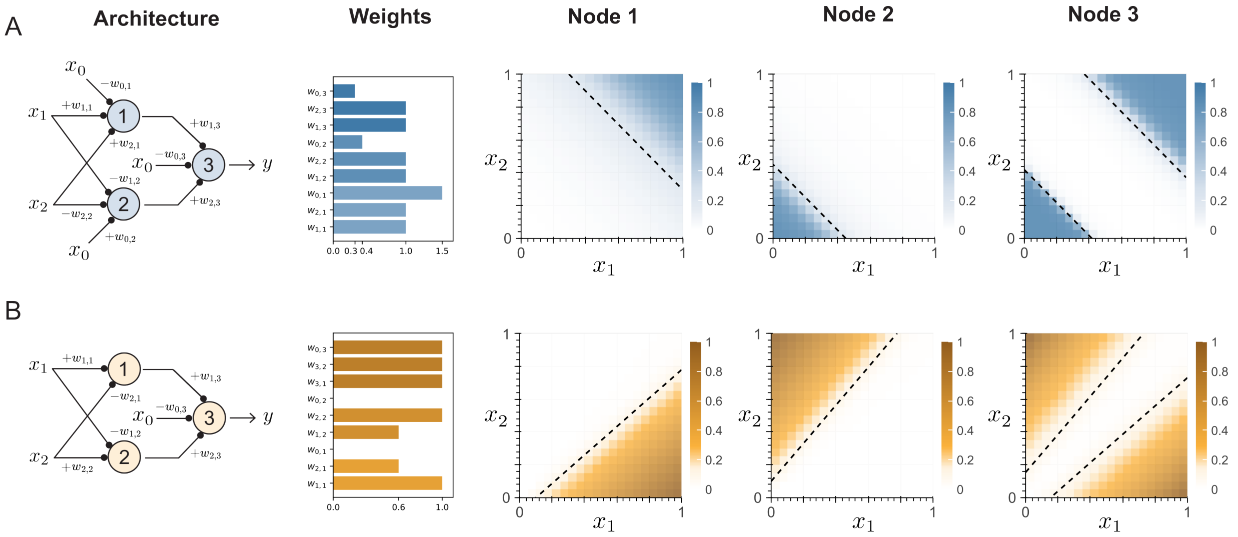

***Figure 3: Design of nonlinear classifiers with a non convex decision boundary*** *Multi-layer perceptrons (MLPs) with 2 nodes in the hidden layer and one node as the output layer were designed based on (A) the activation/deactivation cycle and (B) the catalytic degradation reaction. The sign of each weight (+ or −) was determined from its asymptotic approximation, while the corresponding magnitudes are shown in the bar plots. The absence of a bar indicates that no bias species was included. All heatmaps were normalized by dividing each value by the maximum within each simulation. A grid size of* $N = 20$ *was used for both inputs* $x_1$ *and* $x_2$*, with concentrations ranging from 0 to 1* $\mu$*M. Decision boundaries for each implementation are shown as black dashed lines overlaid on the heatmaps.*

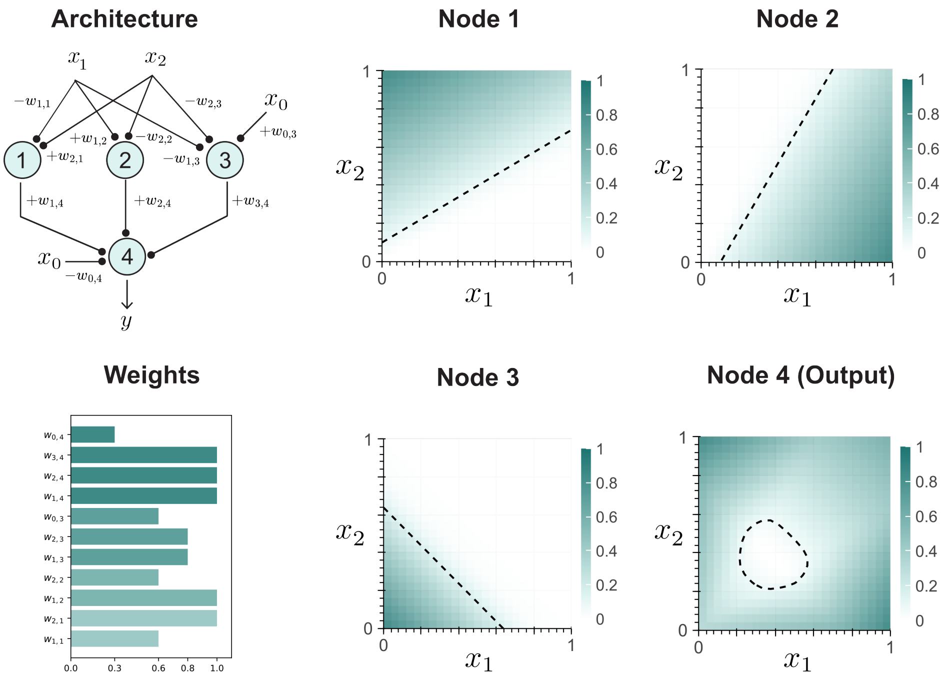

***Figure 4: Design of a sequestration-based nonlinear classifiers with 3 nodes in the hidden layer***. *The sign of each weight (+ or −) was determined from its asymptotic approximation, while the corresponding magnitudes are shown in the bar plots. All heatmaps were normalized by dividing each value by the maximum within each simulation. A grid size of* $N = 20$ *was used for both inputs* $x_1$ *and* $x_2$*, with concentrations ranging from 0 to 1* $\mu$*M. Decision boundaries for each implementation are shown as black dashed lines overlaid on the heatmaps.*

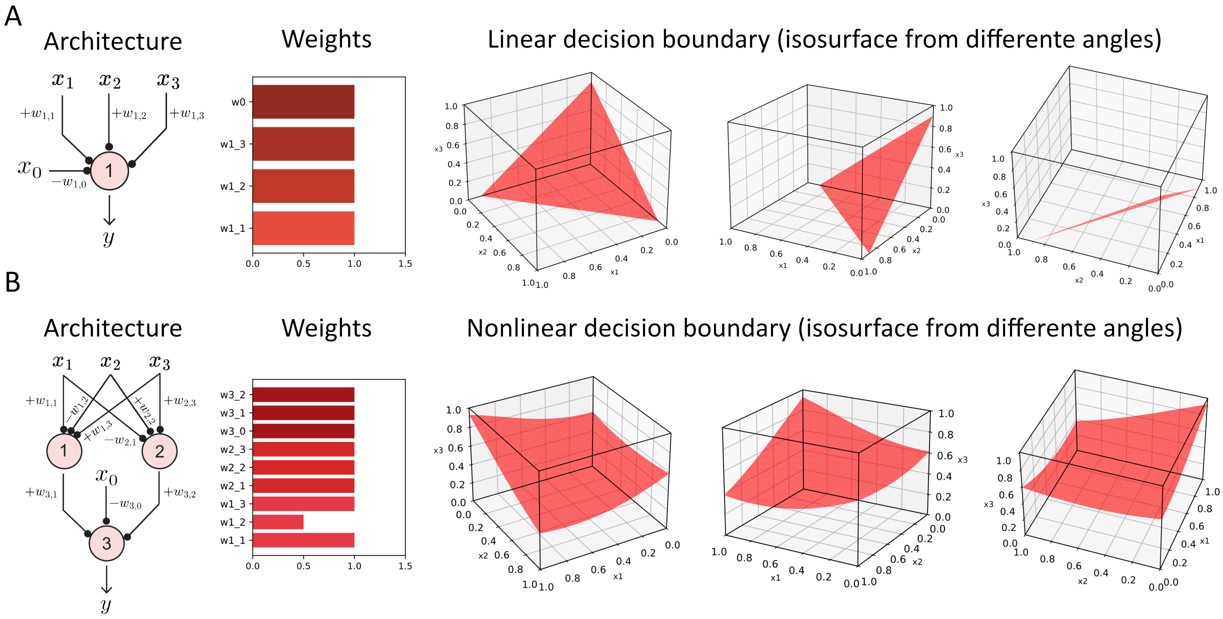

***Figure 5: Design of classifiers with 3 inputs based on the competitive binding mechanism***. *The sign of each weight (+ or −) was determined from its asymptotic approximation, while the corresponding magnitudes are shown in the bar plots. The (A) linear and (B) nonlinear decision boundaries were computed with the isosurface, shown in three different views. A grid size of $N = 20$ was used for all inputs ($x_1$, $x_2$ and $x_3$), with concentrations ranging from 0 to 1 $\mu$M.*

#### 4 Exploratory analysis of the antigen dataset

For the proof-of-concept application of biomolecular neural networks as molecular classifiers, we downloaded the raw dataset from Dannenfelser et al [7] available in their Github repository: `https://github.com/ruthanium/antigen-combos-scripts`, under the name *test-normalized-matrix.txt*. This is a subset of the dataset presented by the authors after batch-correction using COMBAT (`https://rdrr.io/bioc/sva/man/ComBat.html`) and normalization. Expression data is presented in log transformed TPM (transcript per million). To annotate each cancer type to its corresponding tissue, we query both the Cancer Genome Atlas Program (TCGA, `https://www.cancer.gov/ccg/research/genome-sequencing/tcga`) and the Genotype Tissue Expression project (GTEx) [8]. The accession number for each dataset was found in the downloaded dataset, with which we manually annotated based on the clinical data (specifically, pathological reports), available either in the TCGA, or on the original papers that contributed to the dataset. Finally, we assigned each cancer type with their corresponding tissue "label", as shown in Supplementary Table 1.

After annotating the dataset, we set a minimum of 100 samples for either class (cancer or healthy), which resulted in 19 tissue-specific datasets out of 30, as shown in Supplementary Figure 6. To the left of the dashed line, we sorted the datasets by the *minority proportion*, defined as the ratio between the smaller class and the total number of samples. An ideal value of 0.5 suggests a balanced dataset, while lower values indicate stronger representation of one class over the other. In this setting, a larger proportion of healthy cells (blue, in Supplementary Figure 6) may bias classifiers toward false negatives, which are associated with disease recurrence [9, 10, 11]. Conversely, a larger proportion of cancer cells (red, in Supplementary Figure 6) may increase false positives, which in practice could translate into off-target tissue damage [12].

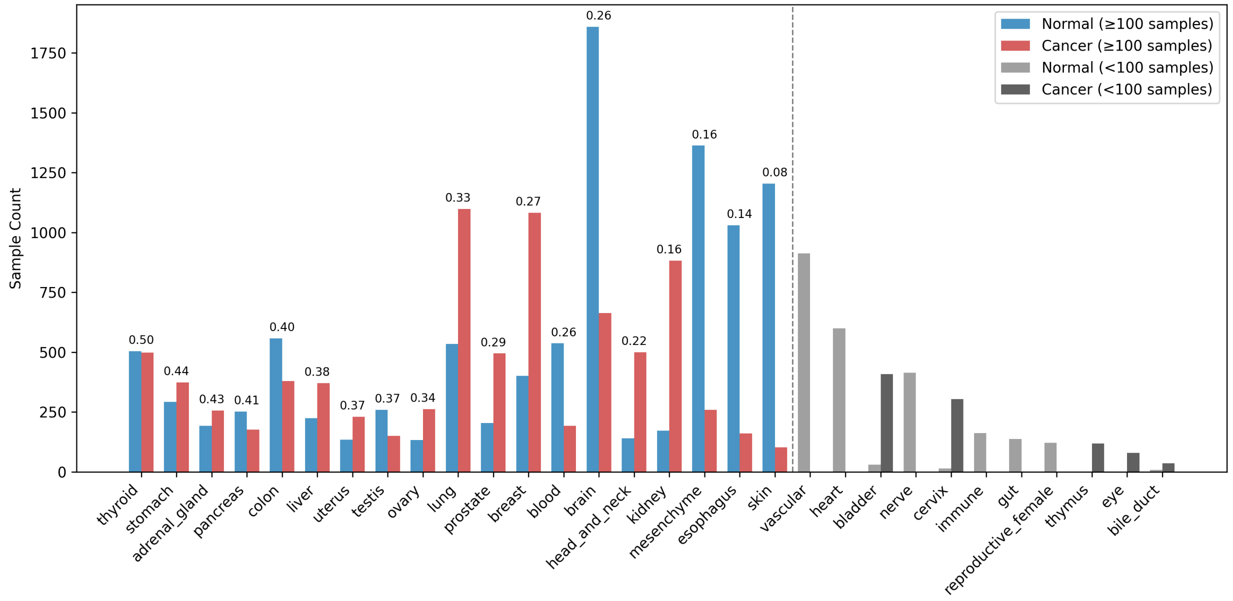

***Figure 6: Class distribution across datasets.*** *Dataset sizes per tissue and class, with cancer samples in red and healthy samples in blue. The minority proportion, computed as the number of samples in the smaller class divided by the total number of samples, is shown above each bar. Tissues are ordered from left to right by minority proportion. Vertical dashed lines separate datasets where either class has fewer than 100 samples.*

To the right of the dashed line in Supplementary Figure 6 are tissues with less than those 100 samples for one of the two classes. For instance, no healthy samples are available for thymus, eye, or bile duct, while no cancer samples are available for gut (small intestine), female reproductive organs (vagina and fallopian tubes), and heart. While the choice of a 100-sample threshold seems arbitrary, a stricter cutoff would have reduced the number of usable datasets to only a few, and thus would not provide a representative case to illustrate the flexibility of our biomolecular neural network design to adapt across different tissue-specific patterns.

For each tissue-specific dataset, we quantified the Earth Mover's Distance (EMD) for every antigen to measure the degree of separation in their expression distributions. Supplementary Figure 7 shows, for each tissue type, the gene with the highest EMD value, which represents the maximum separation observed using a single antigen. Because our dataset is incomplete, we do not interpret antigens with little or no overlap between classes, such as those in testis, ovary, brain, and skin, as novel biomarker candidates. Instead, we reason that these antigens, when combined with others, can contribute to linearly separable combinations. For tissues with highly overlapping expression profiles, such as prostate, blood, kidney, mesenchyme, and esophagus, we interpret classification results with caution, since the degree of overlap limits separability.

We further divided antigens into tertiles based on their EMD values, classifying them into low-, medium-, and high-EMD groups (Supplementary Figure 8). This categorization enabled a systematic exploration of representative expression patterns. We restricted our analysis to pairwise combinations among the ten antigens with the highest EMD values within each category and across categories. This resulted in 45 within-category (considering that $(A, B)$ and $(B, A)$ pairs exhibir the same pattern, and excluding cases such as $(A, A)$) and 100 across-category combinations, covering six EMD group pairs: high–high, high–medium, high–low, medium–medium, medium–low, and low–low.

***Table 1:*** *Cancer and Healthy Tissues' Lookup Table*

| **Original annotation** | **New annotation** |
|---|---|
| Lung | lung |
| Lung Adenocarcinoma | lung |
| Lung Squamous Cell Carcinoma | lung |
| Kidney | kidney |
| Kidney Renal Clear Cell Carcinoma | kidney |
| Kidney Chromophobe | kidney |
| Kidney Renal Papillary Cell Carcinoma | kidney |
| Liver | liver |
| Liver Hepatocellular Carcinoma | liver |
| Bile Duct | bile_duct |
| Cholangiocarcinoma | bile_duct |
| Brain | brain |
| Brain Lower Grade Glioma | brain |
| Glioblastoma Multiforme | brain |
| Prostate | prostate |
| Prostate Adenocarcinoma | prostate |
| Colon | colon |

*Continued on next page*

| Original annotation | New annotation |
|---|---|
| Rectum Adenocarcinoma | colon |
| Colon Adenocarcinoma | colon |
| Skin | skin |
| Skin Cutaneous Melanoma | skin |
| Thymus | thymus |
| Thymoma | thymus |
| Bladder | bladder |
| Bladder Urothelial Carcinoma | bladder |
| Thyroid | thyroid |
| Thyroid Carcinoma | thyroid |
| Pancreas | pancreas |
| Pancreatic Adenocarcinoma | pancreas |
| Breast | breast |
| Breast Invasive Carcinoma | breast |
| Uterus | uterus |
| Uterine Corpus Endometrial Carcinoma | uterus |
| Uterine Carcinosarcoma | uterus |
| Ovary | ovary |
| Ovarian Serous Cystadenocarcinoma | ovary |
| Cervix Uteri | cervix |
| Cervical Squamous Cell Carcinoma and Endocervical Adenocarcinoma | cervix |
| Stomach | stomach |
| Stomach Adenocarcinoma | stomach |
| Esophagus | esophagus |
| Esophageal Carcinoma | esophagus |
| Testis | testis |
| Testicular Germ Cell Tumors | testis |
| Adrenal Gland | adrenal_gland |
| Adrenocortical Carcinoma | adrenal_gland |
| Pheochromocytoma and Paraganglioma | adrenal_gland |
| Blood | blood |
| Acute Myeloid Leukemia | blood |
| Lymphoid Neoplasm Diffuse Large B-cell Lymphoma | blood |
| Mouth | head_and_neck |
| Salivary Gland | head_and_neck |
| Head and Neck Squamous Cell Carcinoma | head_and_neck |
| Nerve | nerve |
| Mesothelioma | lung |
| Muscle | mesenchyme |
| Bone | mesenchyme |
| Adipose Tissue | mesenchyme |
| Sarcoma | mesenchyme |
| Pituitary | brain |
| Vagina | reproductive_female |

*Continued on next page*

| Original annotation | New annotation |
| --- | --- |
| Fallopian Tube | reproductive_female |
| Small Intestine | gut |
| Spleen | immune |
| Heart | heart |
| Blood Vessel | vascular |
| Uveal Melanoma | eye |

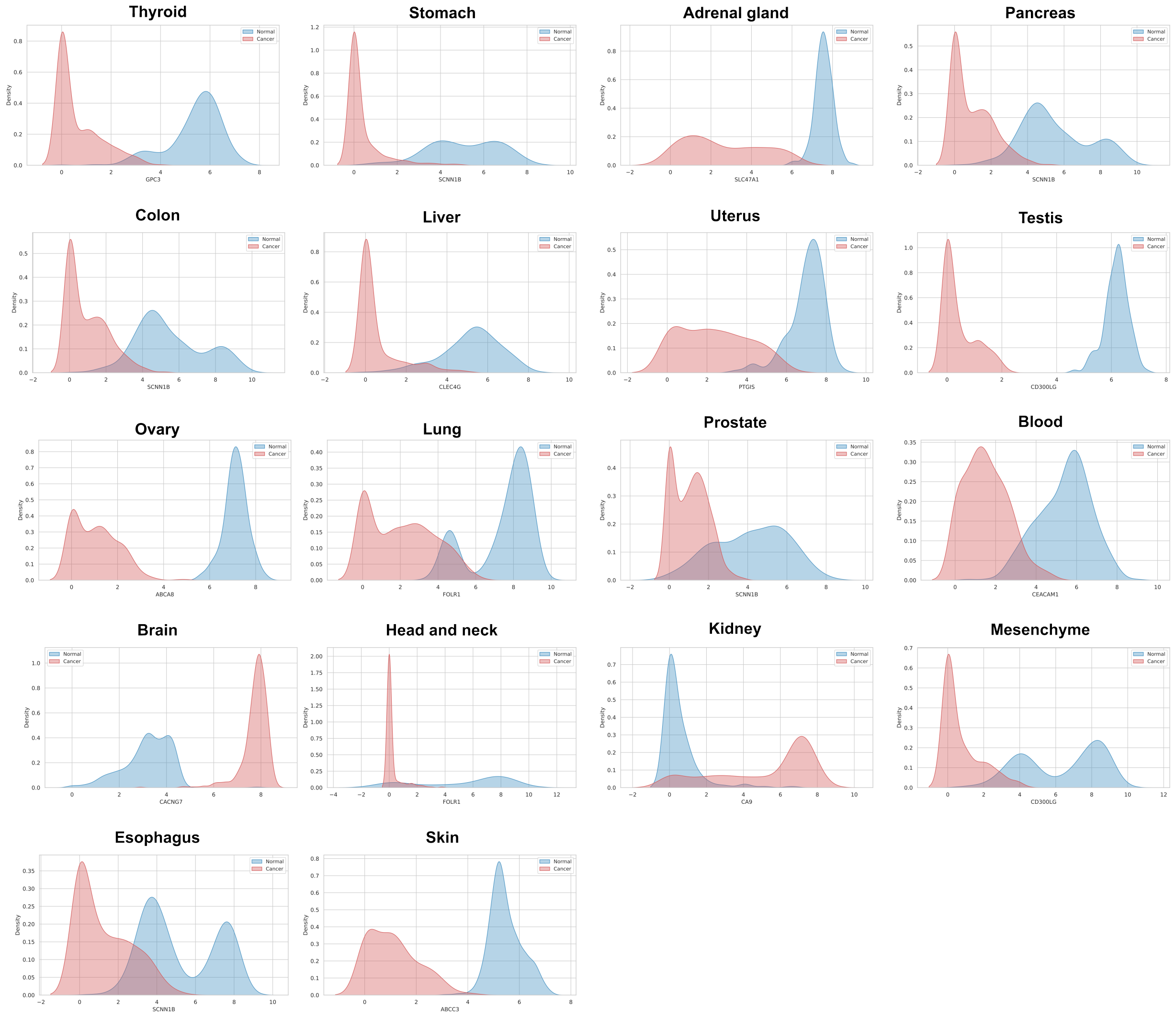

***Figure 7: Distribution of gene expression in cancer and healthy cells for the highest EMD gene per tissue type.*** *Class labels are color-coded: cancer cells in red and healthy cells in blue.*

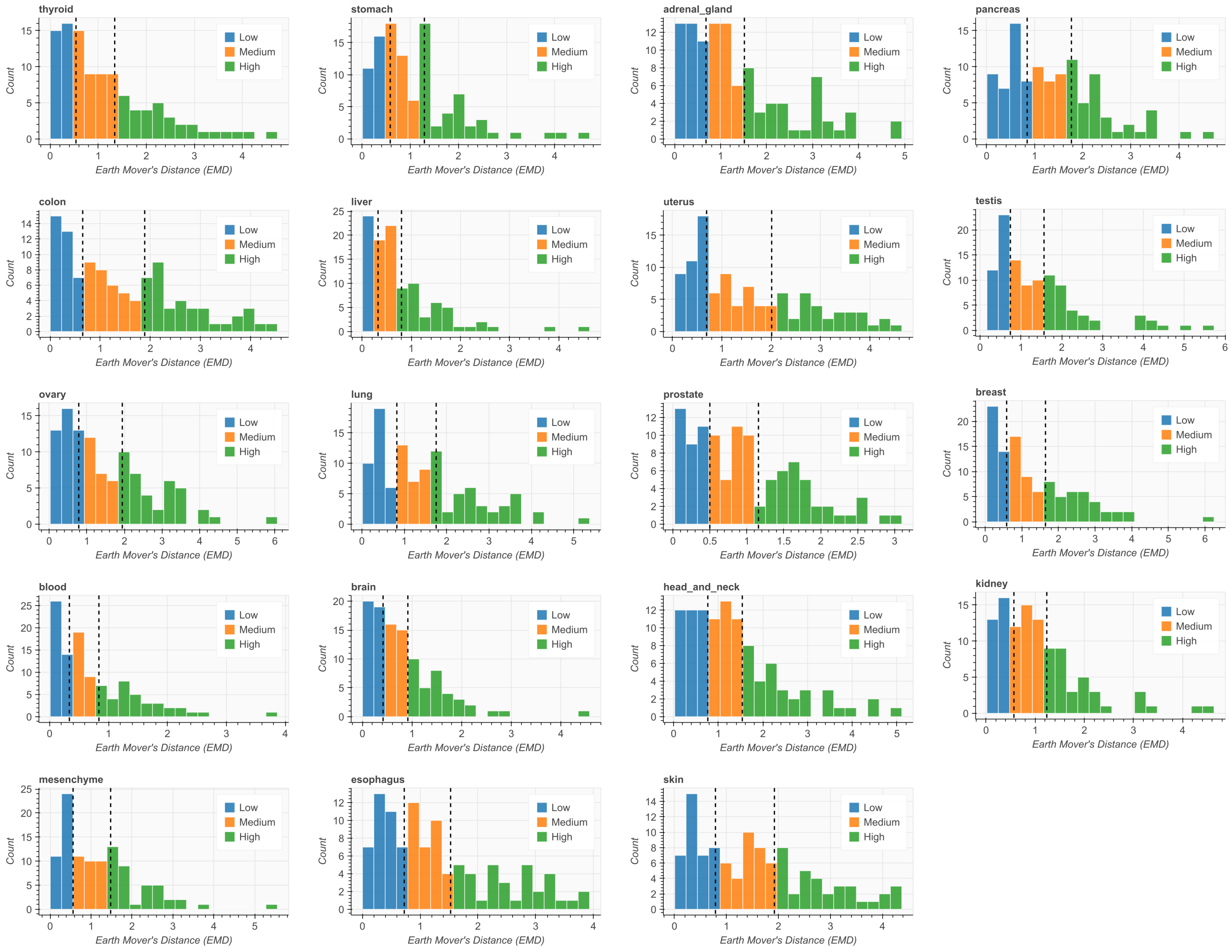

***Figure 8: Class separability measurements using the Earth's Mover Distance (EMD).*** *The black dashed lines indicate the EMD thresholds that define the low, medium, and high EMD categories for each tissue dataset.*

### 5 Design of molecular classifiers for cancer classification

For each within- and across-category combination, we designed linear classifiers based on the activation/deactivation cycle reaction, and nonlinear classifiers with a single hidden layer with 2, 3 or 4 molecular sequestration-based neurons connected to an output layer based on the activation/deactivation cycle reaction. We performed a stratified 5-fold validation to compute the precision, recall, F1 score and area under the ROC curve (AUC) metrics to account for the class imbalance shown in Supplementary Figure 6. The results for each tissue-specific linear classifier are shown in Supplementary Figure 9, whereas the nonlinear classifiers with 2, 3 and 4 nodes in the hidden layer are shown in Supplementary Figures 10 to 12.

#### 5.1 Combinations of high-EMD antigens enable linear separability

We found that including at least one high-EMD antigen consistently results in linear classifiers with high performance. Across all tissues, combinations containing a high-EMD antigen achieved a median F1 score of 0.939 (IQR 0.091), compared to 0.826 (IQR 0.260) for pairs without any high-EMD antigen (computed from Supplementary Figure 9). Within this group, performance decreased in the order High–High, High–Medium, and High–Low, with corresponding medians of 0.939, 0.913, and 0.887, respectively. We confirmed that the presence of at least one antigen that was high-EMD enabled a significant improvement in the F1 score by running a paired Wilcoxon signed-rank test (median difference 0.123, $p = 3.6 \times 10^{-14}$, Cliff's $\delta = 0.63$). Furthermore, to confirm this effect while accounting for inter-tissue variability, we fit a linear mixed-effects model with F1 score as the response variable, tissue as a random intercept, and the presence of a high-EMD antigen as a fixed effect. The model showed that including a high-EMD antigen increased mean F1 by 0.186 (SE = 0.002, $p < 0.001$), indicating that this relationship holds consistently across tissues even when model parameters are shared.

To illustrate these effects, we examined a few antigen pairs that achieved F1, accuracy, recall, and AUC values above 0.9 for each tissue. Supplementary Figures 10–12 show representative expression patterns for the High–High, High–Medium, and High–Low categories, respectively. Interestingly, a small number of Medium–Medium and Medium–Low pairs, particularly in adrenal gland, colon, esophagus, head and neck, mesenchyme, and skin, also achieved performance comparable to high-EMD–containing pairs, as well as Low-Low expression patterns for testis and pancreas (see Supplementary Figure 9), suggesting that tissue-specific expression can occasionally compensate for low antigen discriminatory power. We didn't explore further this observation due to the incompleteness of our dataset.

#### 5.2 Multi-layer perceptrons with increasing width improve performance for medium- and low-EMD antigen combinations

As a strategy to enhance the performance of classifiers for antigen combinations that are not high-EMD, we designed multilayer perceptrons (MLPs) with increasing widths (i.e., more neurons in the hidden layer). Specifically, we compared a linear classifier (no hidden layers) with MLPs containing two (2H), three (3H), and four (4H) hidden layers. To evaluate whether more complex architectures systematically improved classification performance, we first computed pairwise differences in F1 score ($\Delta F_1$) between consecutive models of increasing complexity (Linear→2H, 2H→3H, 3H→4H). We also examined cumulative effects by aggregating the sequential $\Delta F_1$ values across increasing complexity levels (Linear → 2H → 3H and Linear → 2H → 3H → 4H), capturing the total improvement accumulated as additional layers were introduced. These comparisons were then aggregated across the 19 tissues and antigen category pairs (High×High, High×Medium, High×Low, Medium×Medium, Medium×Low, and Low×Low), and shown in Table 2.

For each comparison, we summarized three complementary metrics: (i) the *mean fraction of positive* $\Delta F_1$, representing how often an increase in complexity improved performance; (ii) the *median of medians*, which reports the central tendency of improvement across all tissues; and (iii) the *fraction of tissues showing significant improvement* (FDR < 0.05).

***Table 2:*** *Aggregated improvement in F1 score across biomolecular neural network architectures. Median-of-medians ($\widetilde{\Delta F_1}$), mean fraction of positive $\Delta F_1$, and fraction of tissues showing significant improvement (FDR < 0.05).*

| Comparison | Category pair | Mean fraction positive | Median-of-medians | Fraction tissues significant |
|---|---|---|---|---|
| **Linear → 2H** | | | | |
| | Medium × Low | 0.7932 | 0.0257 | 1.0000 |
| | Medium × Medium | 0.8667 | 0.0257 | 0.8947 |
| | High × Medium | 0.8516 | 0.0207 | 1.0000 |
| | Low × Low | 0.7743 | 0.0182 | 1.0000 |
| | High × High | 0.8865 | 0.0145 | 1.0000 |
| | High × Low | 0.8005 | 0.0116 | 1.0000 |
| **2H → 3H** | | | | |
| | Medium × Medium | 0.6795 | 0.0042 | 0.8947 |
| | Low × Low | 0.5965 | 0.0040 | 0.7368 |
| | Medium × Low | 0.6258 | 0.0026 | 0.8947 |
| | High × Medium | 0.6353 | 0.0015 | 0.8947 |
| | High × Low | 0.6416 | 0.0014 | 0.8947 |
| | High × High | 0.6515 | 0.0011 | 0.8421 |
| **3H → 4H** | | | | |
| | Medium × Low | 0.6321 | 0.0024 | 0.8421 |
| | Medium × Medium | 0.6608 | 0.0022 | 0.7368 |
| | High × Medium | 0.6179 | 0.0014 | 0.7368 |
| | Low × Low | 0.5544 | 0.0012 | 0.4211 |
| | High × Low | 0.5679 | 0.0007 | 0.7895 |
| | High × High | 0.5427 | 0.0005 | 0.6316 |
| **Linear → 3H** | | | | |
| | Medium × Medium | 0.9158 | 0.0301 | 1.0000 |
| | Medium × Low | 0.8547 | 0.0301 | 1.0000 |
| | Low × Low | 0.8000 | 0.0281 | 0.9474 |
| | High × Medium | 0.8968 | 0.0227 | 1.0000 |
| | High × High | 0.9088 | 0.0171 | 1.0000 |
| | High × Low | 0.8605 | 0.0142 | 1.0000 |
| **Linear → 4H** | | | | |
| | Medium × Medium | 0.9287 | 0.0369 | 1.0000 |
| | Low × Low | 0.8140 | 0.0341 | 0.9474 |
| | Medium × Low | 0.8805 | 0.0325 | 1.0000 |
| | High × Medium | 0.9121 | 0.0216 | 1.0000 |
| | High × High | 0.9205 | 0.0172 | 1.0000 |
| | High × Low | 0.8711 | 0.0138 | 1.0000 |

Across all tissues, the largest improvements were observed when moving from the linear model to the 2-hidden-node network (Linear→2H). Median-of-medians ranged from 0.0116 to 0.0257 across category pairs, with nearly all tissues showing significant gains (FDR < 0.05). Adding additional nodes led to smaller average improvements: for 2H→3H, typical median-of-medians were below 0.004, and for 3H→4H, below 0.002. Even so, 60–80% of antigen pairs still showed positive $\Delta F_1$ when increasing the number of nodes in the hidden layer. The cumulative comparisons (Linear→2H→3H and Linear→2H→3H→4H) showed total median gains around 0.02–0.04, confirming that most of the improvement occurs in the initial transition from a linear to a nonlinear classifier. Medium×Medium and Medium×Low pairs benefited the most from additional nodes, while Low×Low pairs consistently showed the smallest and least consistent improvements. Furthermore, since we observe diminishing "advantage" when increasing the network complexity, and given that the F1 scores were computed using a 5-fold stratified cross-validation, we consider that none of the results shown throughout Supplementary Figure 9 to 12 are cases of overfitting.

To evaluate whether the slight improvements observed in the aggregate results were consistent within individual tissues, we examined per-tissue $\Delta F_1$ distributions for each model comparison (Supplementary Figures 16–20). In these plots, the $x$-axis corresponds to antigen category pairs and the $y$-axis to the change in F1 score ($\Delta F_1$). Consistent with the aggregate analysis, increasing model complexity generally led to more antigen pairs with positive $\Delta F_1$ values, visible as a larger proportion of gray points above zero. The top five antigen pairs with the largest performance increases are highlighted in each panel. Interestingly, although many antigen combinations benefited from more nodes in the hidden layers, pairs within the Low$\times$Low group remained poorly separable regardless of architecture. Ultimately, Supplementary Figure 21 shows representative decision boundaries for the 2H, 3H, and 4H models, highlighting the models' expressiveness across most antigen categories.

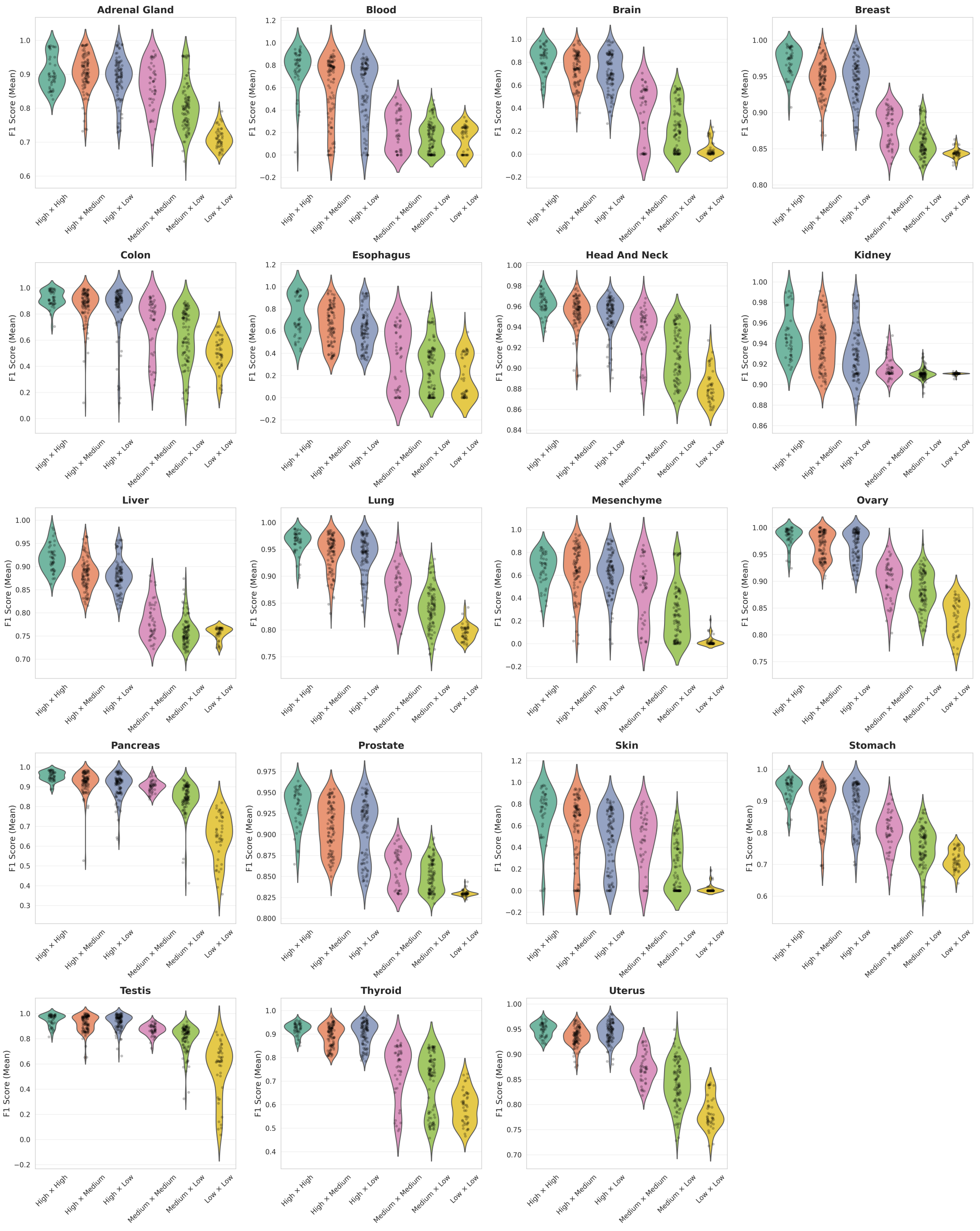

***Figure 9: F1 score distributions across EMD-defined groups of tissue-specific linear classifiers.*** *Each dot in the violin plots represents one antigen pair whose EMD value falls within the EMD range indicated on the x-axis. Distributions for within-category comparisons (High–High, Medium–Medium, and Low–Low) include* $n = 45$ *unique antigen pairs (55 were duplicate combinations). Among-category comparisons include* $n = 100$ *antigen pairs per group. The F1 score was computed as the mean across five cross-validation folds. The standard deviation per antigen combination is not shown.*

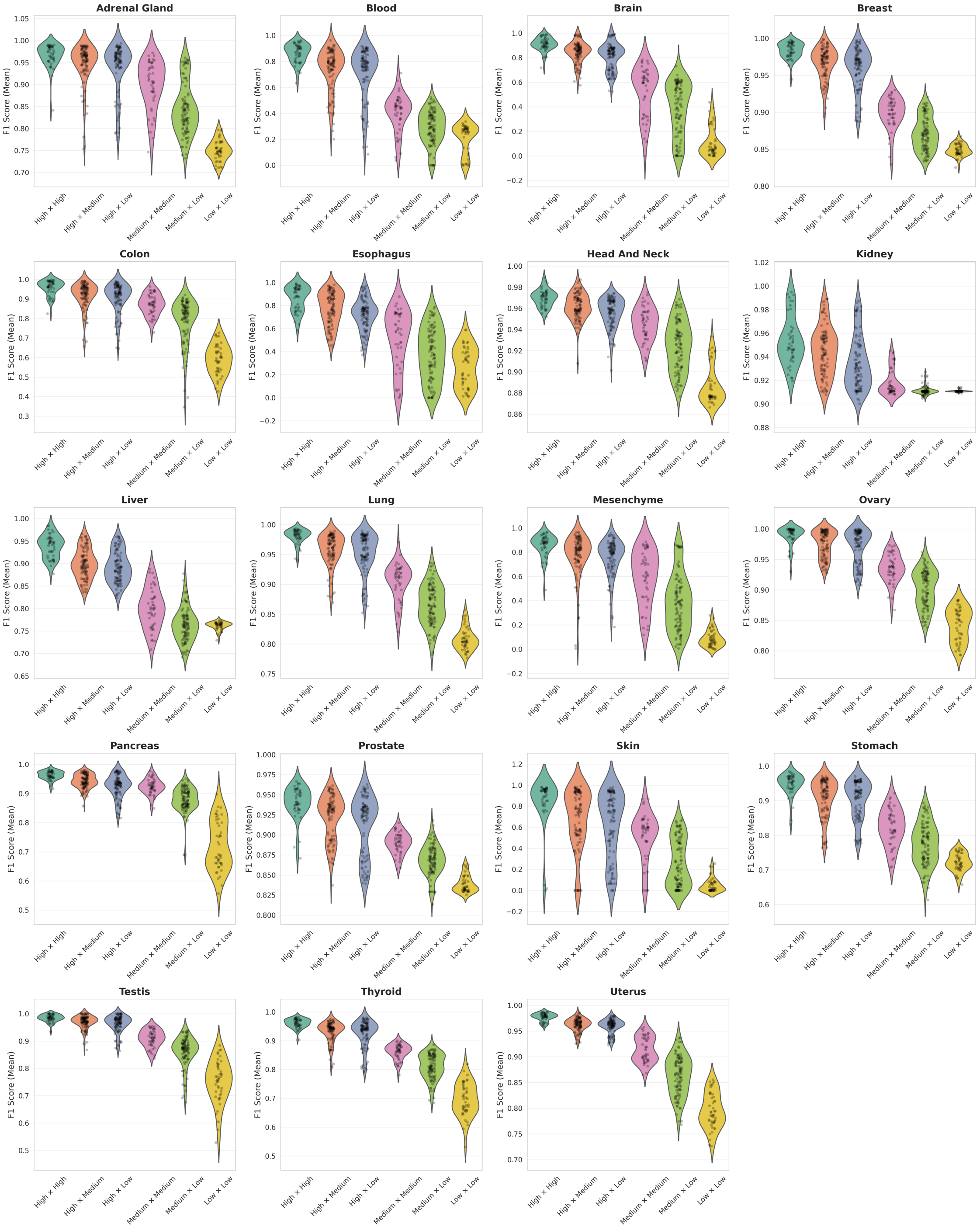

***Figure 10: F1 score distributions across EMD-defined groups of tissue-specific nonlinear classifiers buildt using 2 nodes in the hidden layer.*** *Each dot in the violin plots represents one antigen pair whose EMD value falls within the EMD range indicated on the x-axis. Distributions for within-category comparisons (High–High, Medium–Medium, and Low–Low) include* $n = 45$ *unique antigen pairs (55 were duplicate combinations). Among-category comparisons include* $n = 100$ *antigen pairs per group. The F1 score was computed as the mean across five cross-validation folds. The standard deviation per antigen combination is not shown.*

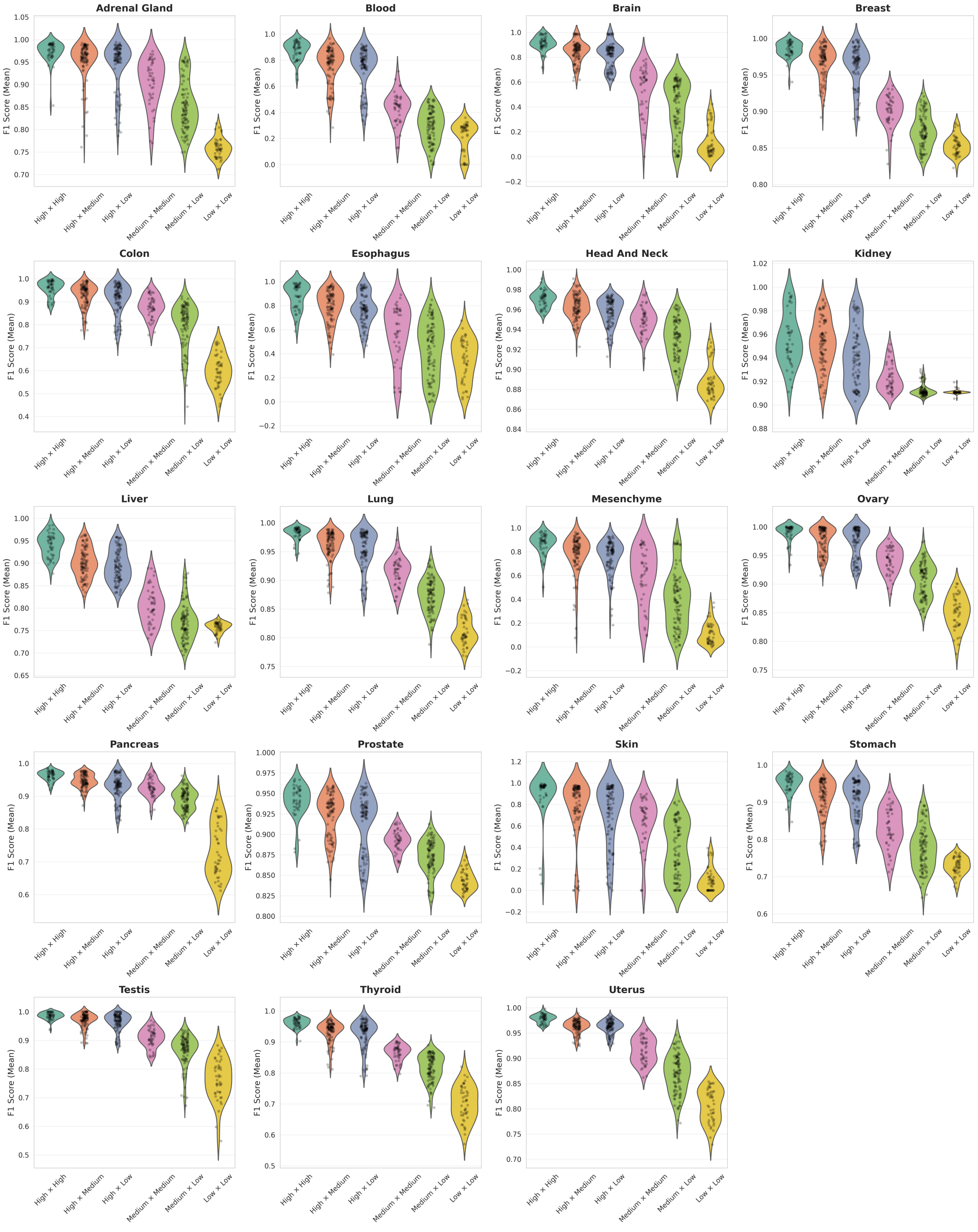

***Figure 11: F1 score distributions across EMD-defined groups of tissue-specific nonlinear classifiers buildt using 3 nodes in the hidden layer.*** *Each dot in the violin plots represents one antigen pair whose EMD value falls within the EMD range indicated on the x-axis. Distributions for within-category comparisons (High–High, Medium–Medium, and Low–Low) include* $n = 45$ *unique antigen pairs (55 were duplicate combinations). Among-category comparisons include* $n = 100$ *antigen pairs per group. The F1 score was computed as the mean across five cross-validation folds. The standard deviation per antigen combination is not shown.*

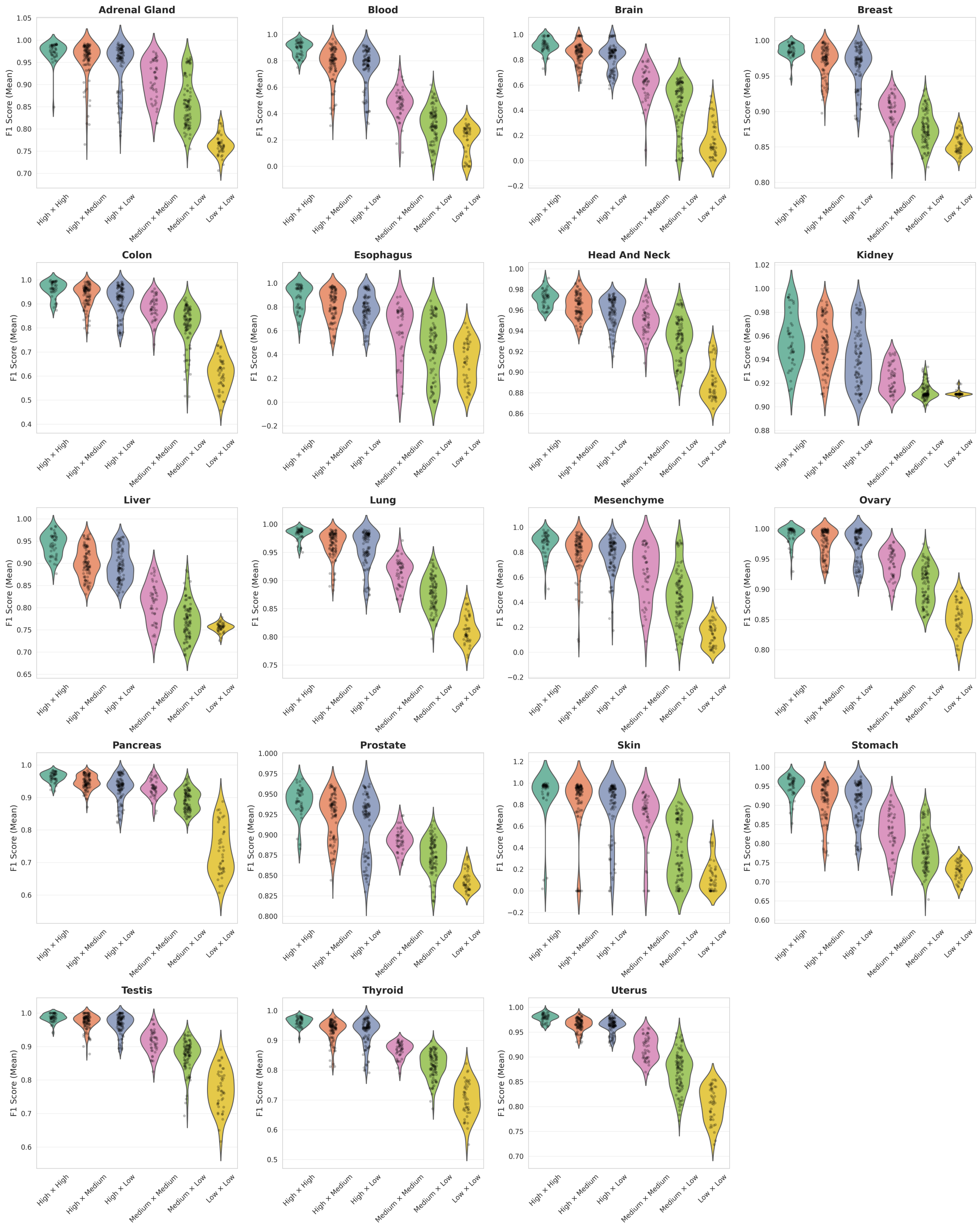

***Figure 12: F1 score distributions across EMD-defined groups of tissue-specific nonlinear classifiers buildt using 4 nodes in the hidden layer.*** *Each dot in the violin plots represents one antigen pair whose EMD value falls within the EMD range indicated on the x-axis. Distributions for within-category comparisons (High–High, Medium–Medium, and Low–Low) include* $n = 45$ *unique antigen pairs (55 were duplicate combinations). Among-category comparisons include* $n = 100$ *antigen pairs per group. The F1 score was computed as the mean across five cross-validation folds. The standard deviation per antigen combination is not shown.*

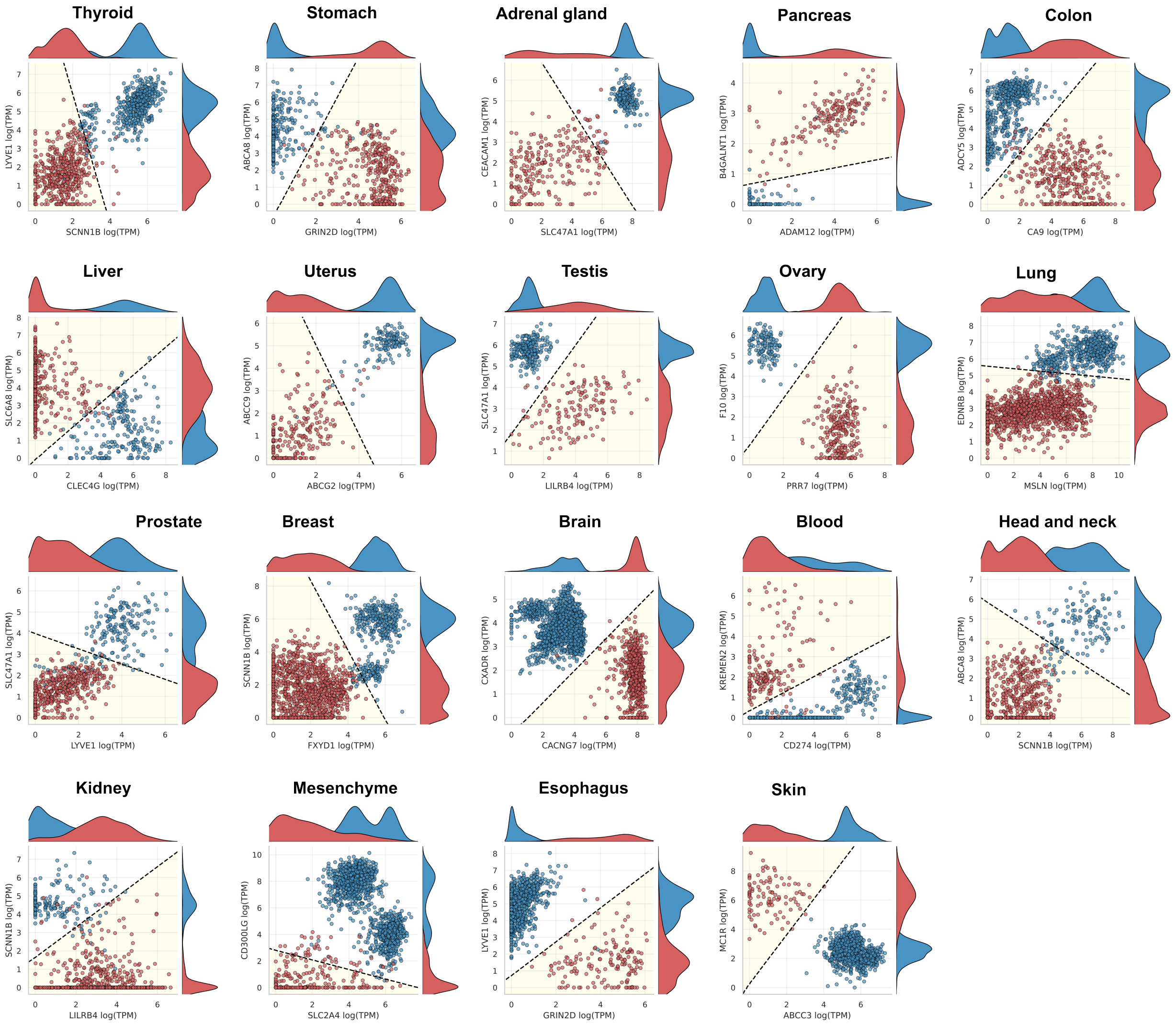

***Figure 13: Tissue-specific linear classifiers when both antigens exhibit high EMD.*** *The linear decision boundary of each classifier is shown as a dashed black line, with the region classified as cancer highlighted in yellow. True sample labels are color-coded, with cancer samples in red and healthy samples in blue. Performance metrics (precision, recall, F1 score, and AUC) were computed across five cross-validation folds, all achieving values of 0.9 or higher.*

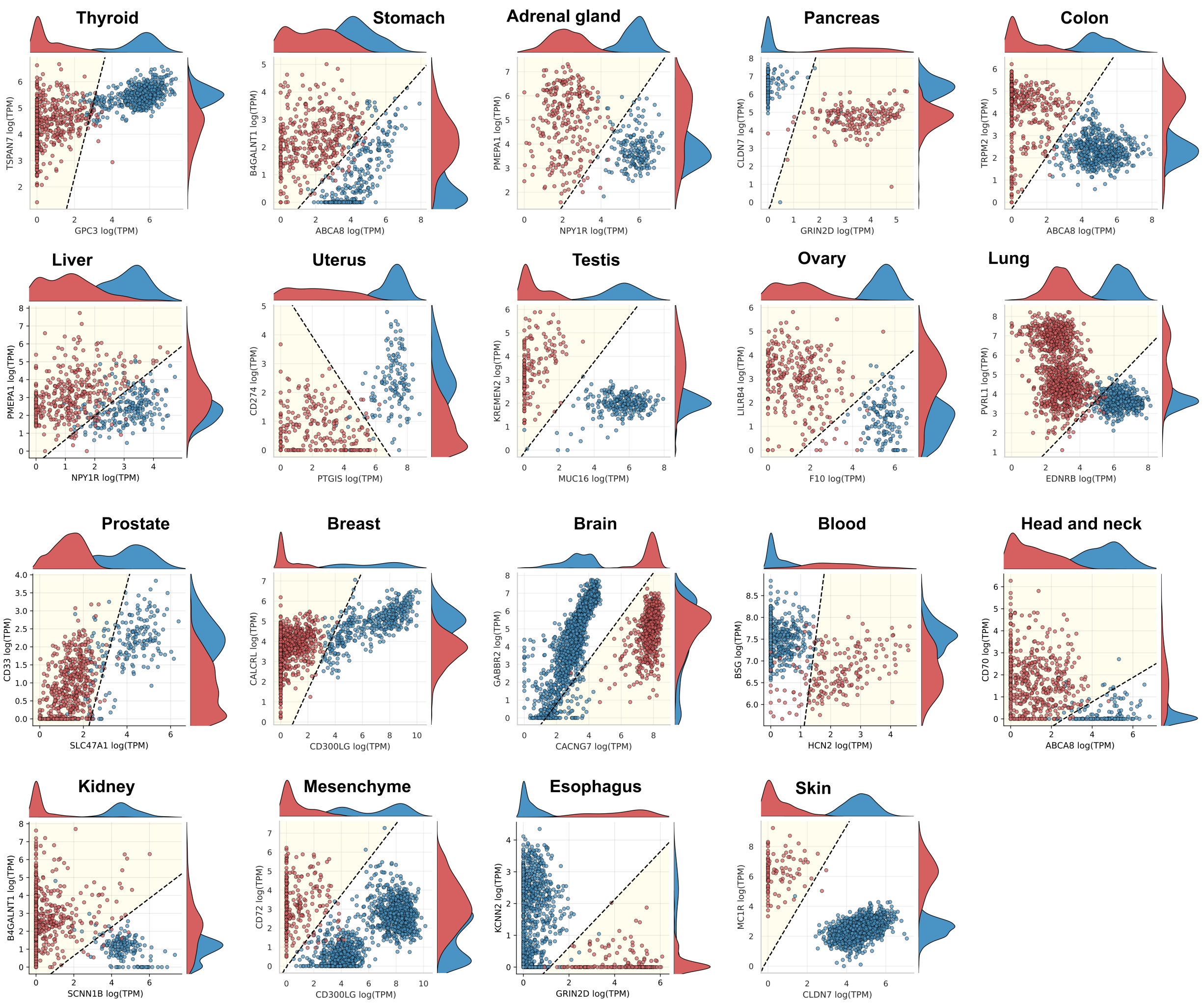

***Figure 14: Tissue-specific linear classifiers when one antigen exhibits high EMD, while the other medium EMD.*** *The linear decision boundary of each classifier is shown as a dashed black line, with the region classified as cancer highlighted in yellow. True sample labels are color-coded, with cancer samples in red and healthy samples in blue. Performance metrics (precision, recall, F1 score, and AUC) were computed across five cross-validation folds, all achieving values of 0.9 or higher.*

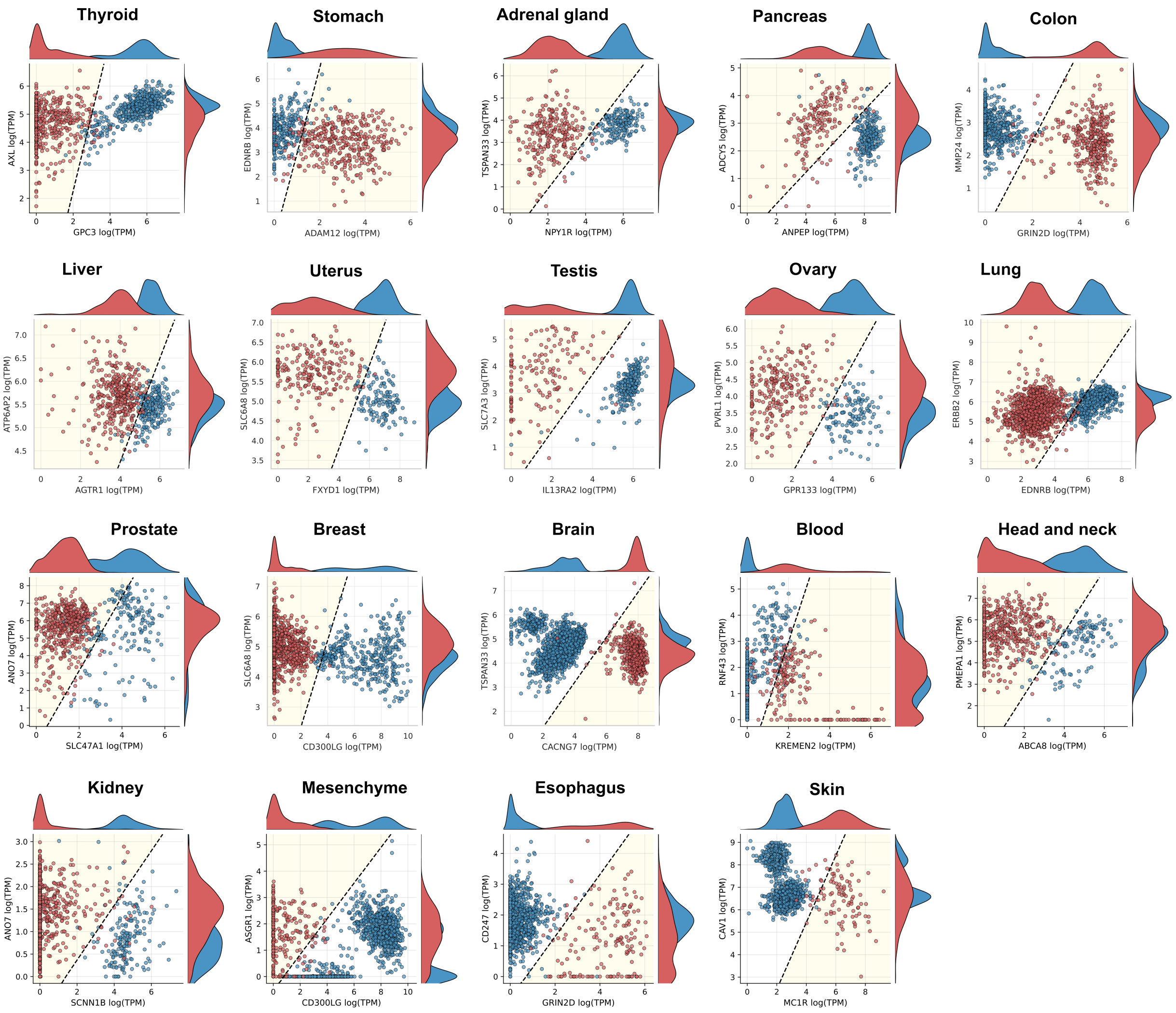

***Figure 15: Tissue-specific linear classifiers when one antigen exhibits high EMD, while the other low EMD.*** *The linear decision boundary of each classifier is shown as a dashed black line, with the region classified as cancer highlighted in yellow. True sample labels are color-coded, with cancer samples in red and healthy samples in blue. Performance metrics (precision, recall, F1 score, and AUC) were computed across five cross-validation folds, all achieving values of 0.9 or higher.*

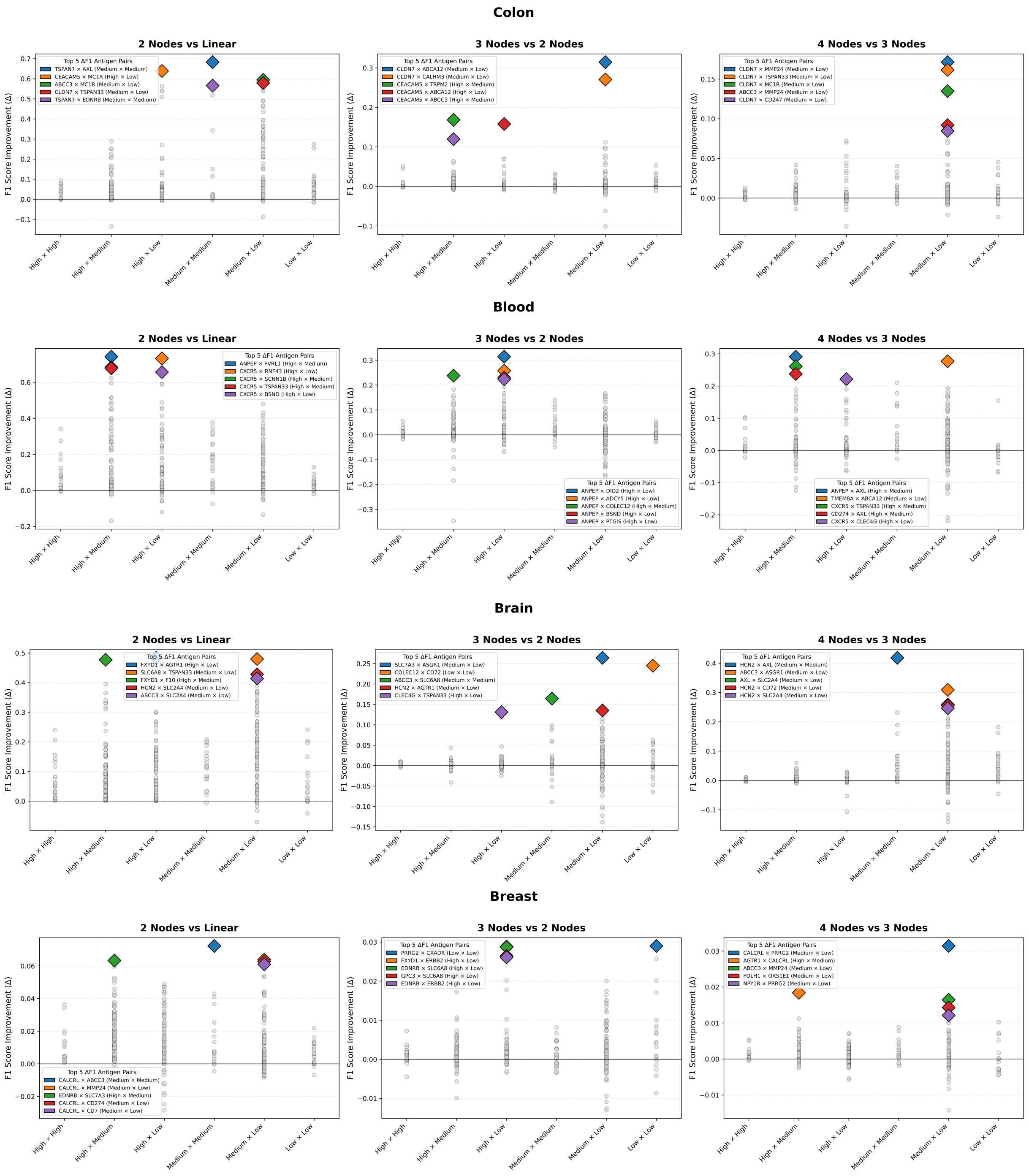

***Figure 16: F1 score improvement among neural networks architectures designed for tcolon, blood, brain and breast tissue.*** *Each panel compares the performance between two specific network architectures: linear classifiers versus 2-node networks (left), 2-node versus 3-node networks (center), and 3-node versus 4-node networks (right). The y-axis represents the change in F1 score (ΔF1), calculated as the difference between the more complex and simpler architecture for each antigen pair. Gray circles represent all antigen pair comparisons, grouped by expression category combinations (High, Medium, Low). Colored diamonds highlight the top 5 antigens that show the highest performance increase upon increasing the network's complexity*

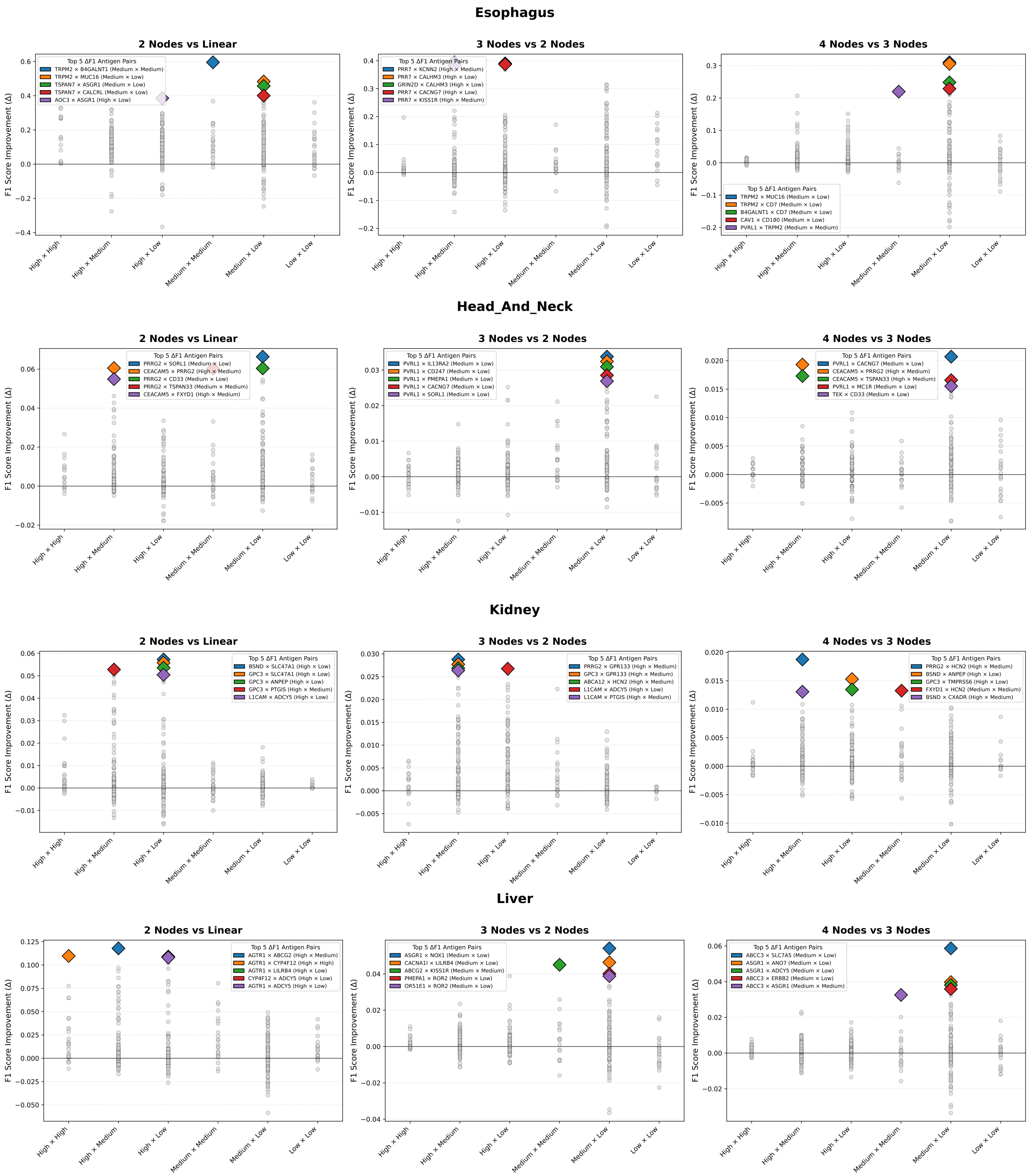

***Figure 17: F1 score improvement among neural networks architectures designed for esophagus, head and neck, kidney, and liver tissue.*** *Each panel compares the performance between two specific network architectures: linear classifiers versus 2-node networks (left), 2-node versus 3-node networks (center), and 3-node versus 4-node networks (right). The y-axis represents the change in F1 score (*$\Delta$*F1), calculated as the difference between the more complex and simpler architecture for each antigen pair. Gray circles represent all antigen pair comparisons, grouped by expression category combinations (High, Medium, Low). Colored diamonds highlight the top 5 antigens that show the highest performance increase upon increasing the network's complexity*

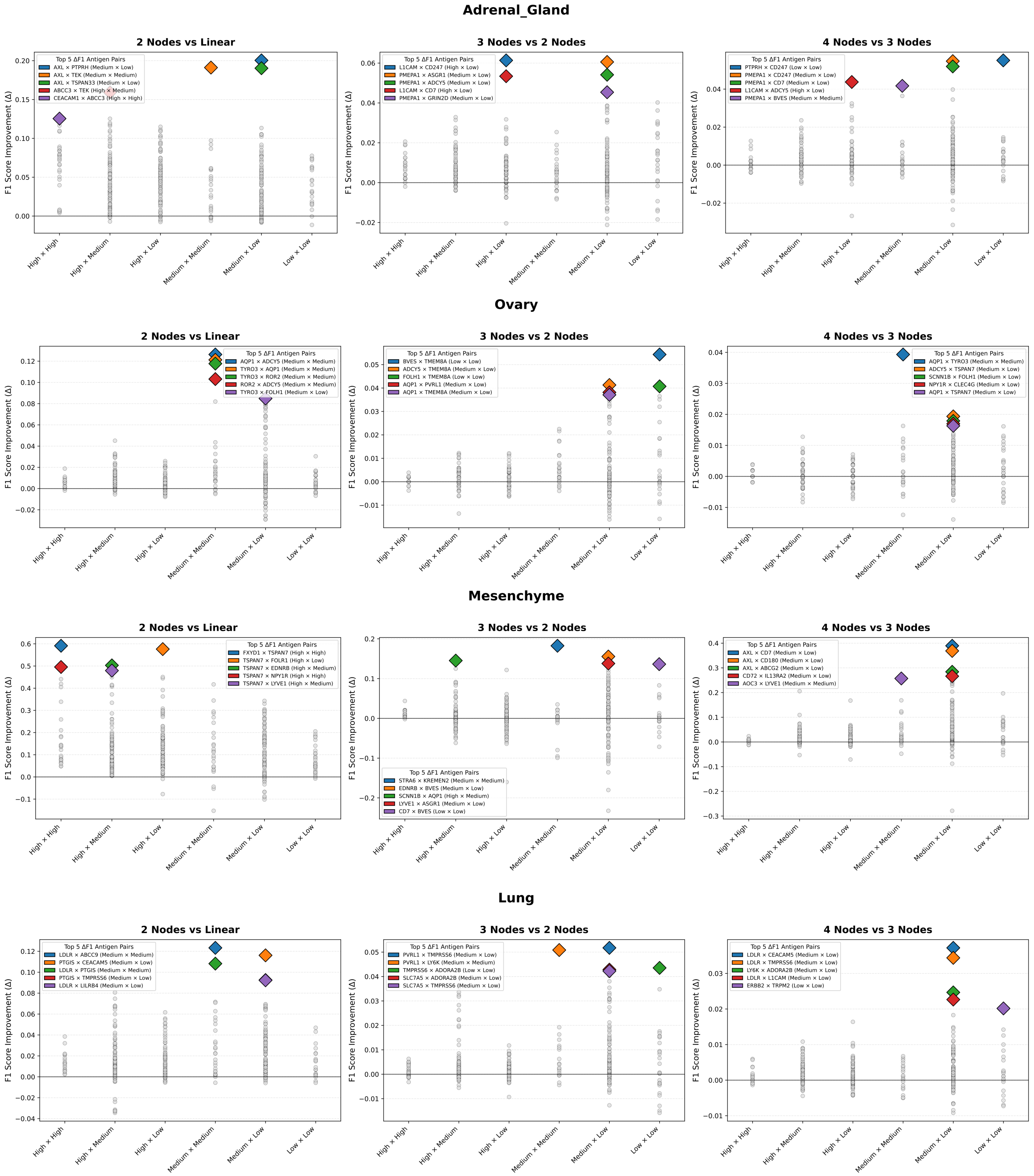

***Figure 18: F1 score improvement among neural networks architectures designed for adrenal gland, ovary, mesenchyme, and lung tissue.*** *Each panel compares the performance between two specific network architectures: linear classifiers versus 2-node networks (left), 2-node versus 3-node networks (center), and 3-node versus 4-node networks (right). The y-axis represents the change in F1 score (ΔF1), calculated as the difference between the more complex and simpler architecture for each antigen pair. Gray circles represent all antigen pair comparisons, grouped by expression category combinations (High, Medium, Low). Colored diamonds highlight the top 5 antigens that show the highest performance increase upon increasing the network's complexity*

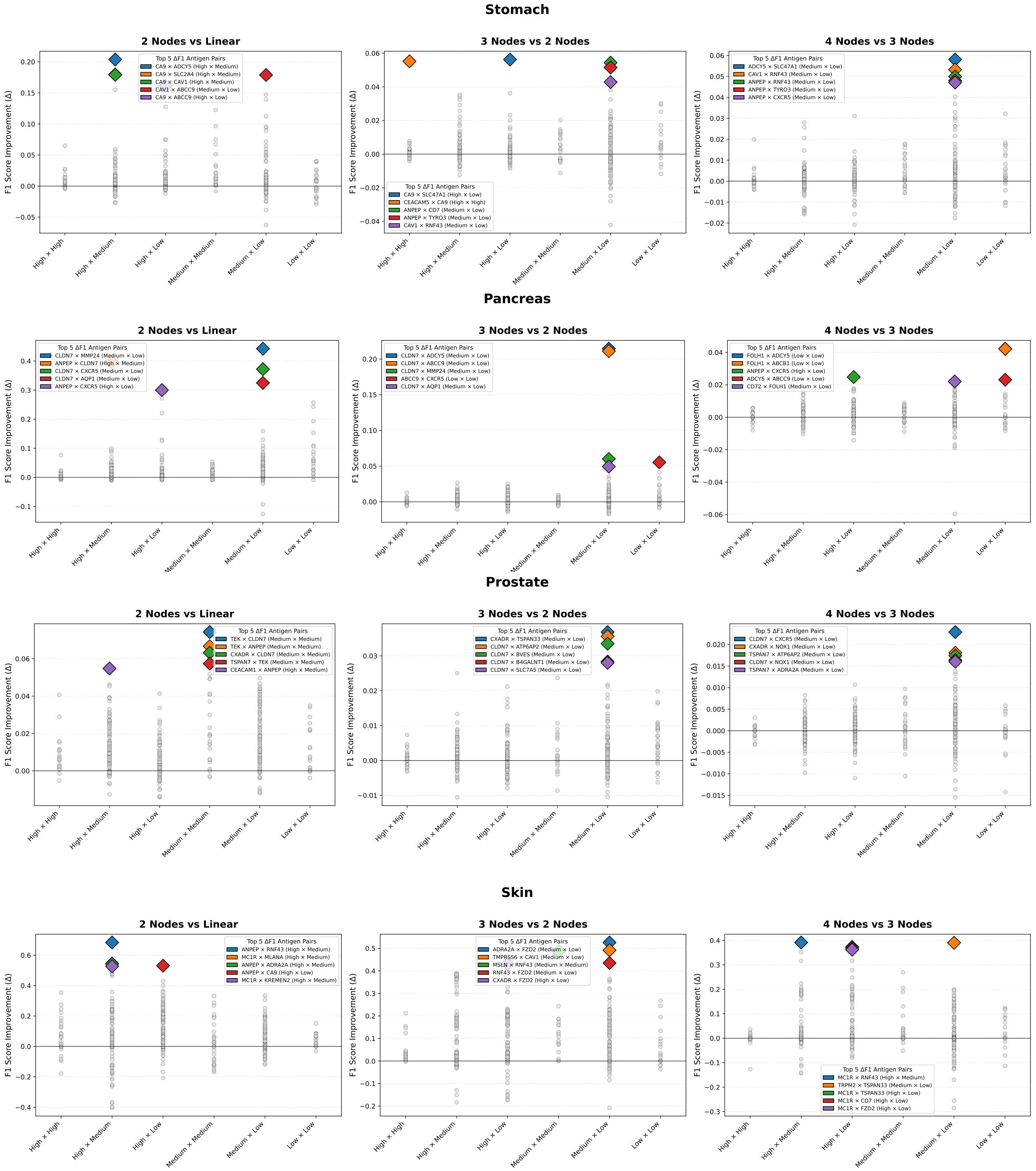

***Figure 19: F1 score improvement among neural networks architectures designed for stomach, pancreas, prostate, and skin tissue.*** *Each panel compares the performance between two specific network architectures: linear classifiers versus 2-node networks (left), 2-node versus 3-node networks (center), and 3-node versus 4-node networks (right). The y-axis represents the change in F1 score ($\Delta F1$), calculated as the difference between the more complex and simpler architecture for each antigen pair. Gray circles represent all antigen pair comparisons, grouped by expression category combinations (High, Medium, Low). Colored diamonds highlight the top 5 antigens that show the highest performance increase upon increasing the network's complexity*

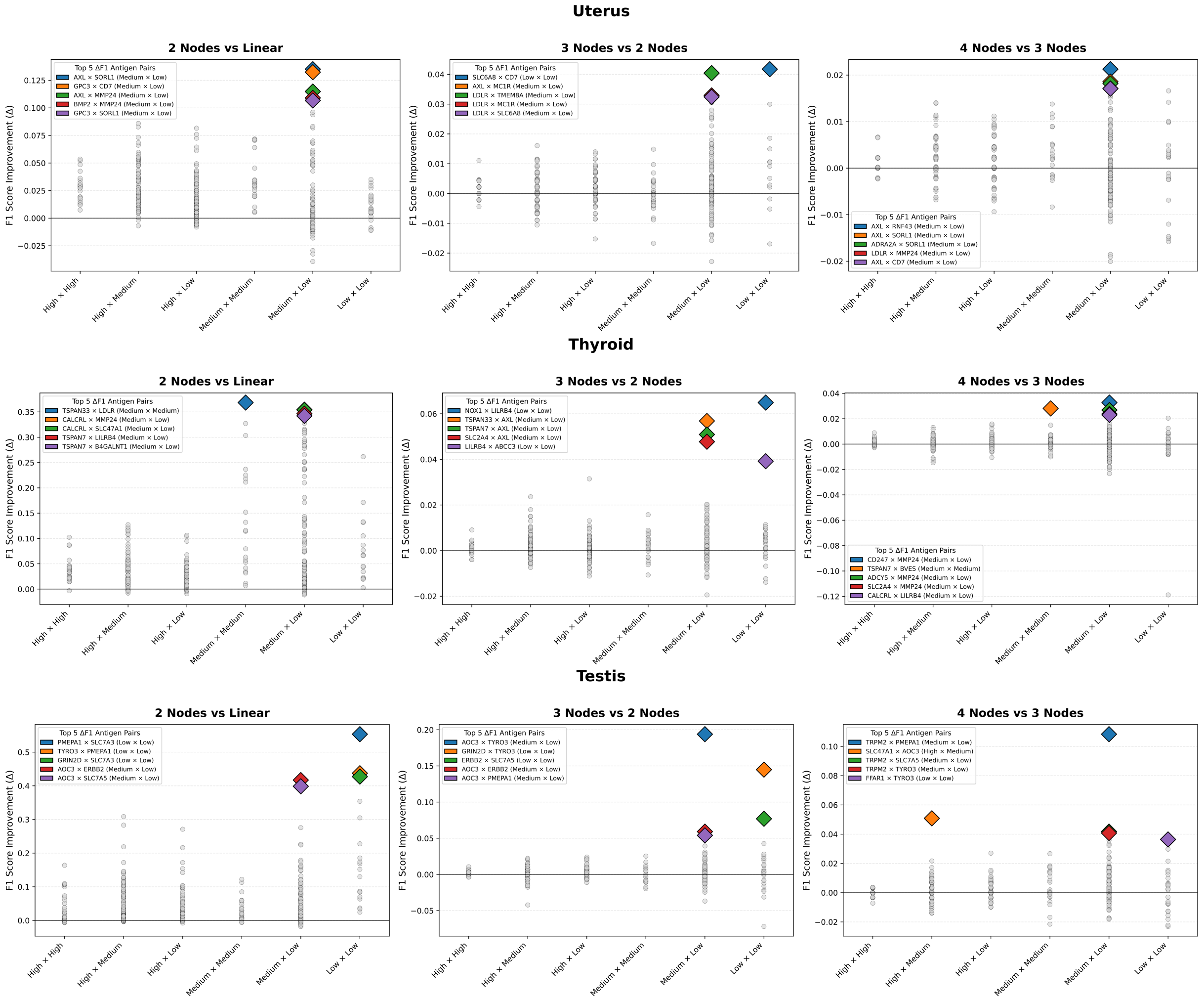

***Figure 20: F1 score improvement among neural networks architectures designed for uterus, thyroid and testis tissue.*** *Each panel compares the performance between two specific network architectures: linear classifiers versus 2-node networks (left), 2-node versus 3-node networks (center), and 3-node versus 4-node networks (right). The y-axis represents the change in F1 score (ΔF1), calculated as the difference between the more complex and simpler architecture for each antigen pair. Gray circles represent all antigen pair comparisons, grouped by expression category combinations (High, Medium, Low). Colored diamonds highlight the top 5 antigens that show the highest performance increase upon increasing the network's complexity*

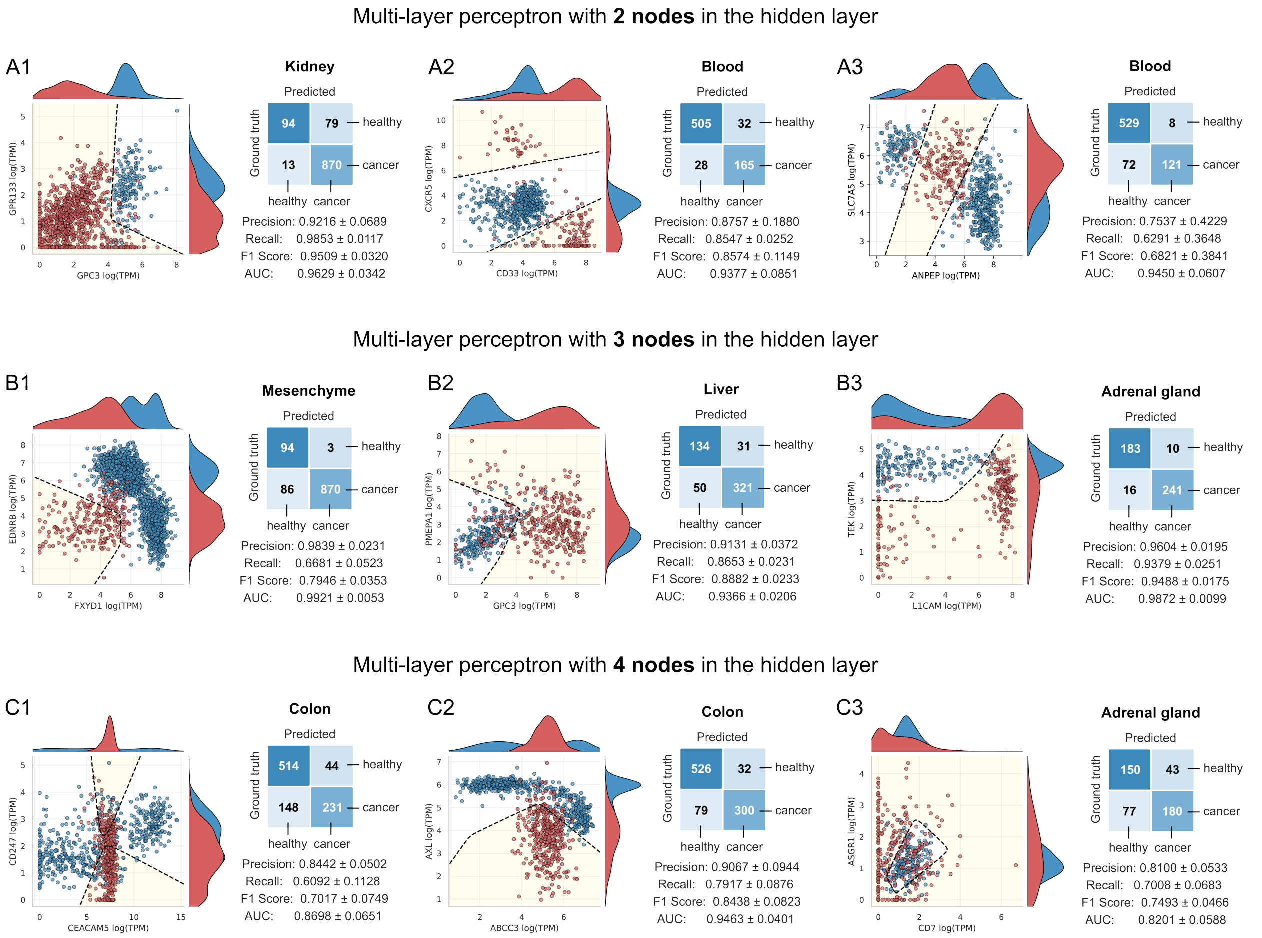

***Figure 21: Some examples of tissue-specific nonlinear classifiers.*** *The decision boundary for each architecture is shown as a dashed black line, corresponding to networks with (A1–A3) 2, (B1–B3) 3, and (C1–C3) 4 hidden nodes, each connected to a single output node. The region classified as cancer is highlighted in yellow. True sample labels are color-coded, with cancer samples in red and healthy samples in blue. To the right of each pattern plot, the aggregate confusion matrix from 5-fold cross-validation is shown. Performance metrics represent the mean ± standard deviation across the five folds.*

[11] Ana Alarcon Tomas, Joshua A. Fein, Shalev Fried, Jessica R. Flynn, Sean M. Devlin, Warren B. Fingrut, Theodora Anagnostou, Anna Alperovich, Nishi Shah, Ellen Fraint, Richard J. Lin, Michael Scordo, Connie Lee Batlevi, Michal J. Besser, Parastoo B. Dahi, Ivetta Danylesko, Sergio Giralt, Brandon S. Imber, Elad Jacoby, Meirav Kedmi, Arnon Nagler, M. Lia Palomba, Mikhail Roshal, Gilles A. Salles, Craig Sauter, Noga Shem-Tov, Avichai Shimoni, Joachim Yahalom, Ronit Yerushalmi, Gunjan L. Shah, Abraham Avigdor, Miguel-Angel Perales, and Roni Shouval. Outcomes of first therapy after cd19-car-t treatment failure in large b-cell lymphoma. *Leukemia*, 37(1):154–163, November 2022.

[12] Gideon Gross and Zelig Eshhar. Therapeutic potential of t cell chimeric antigen receptors (cars) in cancer treatment: Counteracting off-tumor toxicities for safe car t cell therapy. *Annual Review of Pharmacology and Toxicology*, 56(1):59–83, January 2016.
